# COMPARISON OF HIGH-THROUGHPUT SINGLE-CELL RNA-SEQ METHODS FOR EX VIVO DRUG SCREENING

**DOI:** 10.1101/2023.09.28.560069

**Authors:** Henrik Gezelius, Anna Pia Enblad, Anders Lundmark, Martin Åberg, Kristin Blom, Jakob Rudfeldt, Amanda Raine, Arja Harila, Verónica Rendo, Merja Heinäniemi, Claes Andersson, Jessica Nordlund

**Affiliations:** Department of Medical Sciences and Science for Life Laboratory, Uppsala University, Uppsala, 751 44, Sweden; Department of Women’s and Children’s Health, Uppsala University, Uppsala, 751 85, Sweden; Department of Clinical Chemistry and Pharmacology, Uppsala University Hospital, Uppsala, 751 85, Sweden; Department of Immunology, Genetics and Pathology, Uppsala University, Uppsala, 751 85, Sweden; School of Medicine, University of Eastern Finland, 70210 Kuopio, Finland

## Abstract

Functional precision medicine (FPM) aims to optimize patient-specific drug selection based on the unique characteristics of their cancer cells. Recent advancements in high throughput *ex vivo* drug profiling have accelerated interest in FPM. Here, we present a proof-of-concept study for an integrated experimental system that incorporates *ex vivo* treatment response with a single-cell gene expression output enabling barcoding of several drug conditions in one single-cell sequencing experiment. We demonstrate this through a proof-of-concept investigation focusing on the glucocorticoid-resistant acute lymphoblastic leukemia (ALL) E/R+ Reh cell line. Three different single-cell transcriptome sequencing (scRNA-seq) approaches were evaluated, each exhibiting high cell recovery and accurate tagging of distinct drug conditions. Notably, our comprehensive analysis revealed variations in library complexity, sensitivity (gene detection), and differential gene expression detection across the methods. Despite these differences, we identified a substantial transcriptional response to fludarabine, a highly relevant drug for treating high-risk ALL, which was consistently recapitulated by all three methods. These findings highlight the potential of our integrated approach for studying drug responses at the single-cell level and emphasize the importance of method selection in scRNA-seq studies. Finally, our data encompassing 27,327 cells are freely available to extend to future scRNA-seq methodological comparisons.

## INTRODUCTION

Functional precision medicine (FPM) is a strategy whereby live cancer cells are perturbed with drugs, *ex vivo*, which has translational potential to provide personalized information for guiding therapy *in vivo* (1). The model of applying a drug to cancer cells and observing its behavior is not a new concept (2). Early initiatives towards clinical translation faced substantial problems, including poor growth of cells in *ex vivo* cultures, low throughput culture conditions, and limited availability of new drugs to analyze. The past decade yielded dramatic increases in new high throughput technologies and a dramatic increase in the number of drugs available, as well as widespread adoption of comprehensive genomic analysis of cancer samples (3–7). This has dramatically revived interest in *ex vivo* analysis as a method to guide clinical treatment (8–10).

In parallel, single-cell sequencing has greatly improved our understanding of molecular heterogeneity in cancer (11, 12). In the context of FPM, identifying the genes and pathways underlying the heterogeneous behaviors of individual cells in response to a specific treatment is an important step to improve biological understanding and efficacy of treatment (13). Combining FPM with single-cell output for investigating cell-specific *ex vivo* treatment response holds great promise for understanding cellular responses to drugs that may be overlooked in bulk assays (14, 15). While *ex vivo* drug screening has advanced our understanding of cellular responses to drugs, significant challenges remain with incorporating single-cell RNA-seq (scRNA-seq) as an output. One major difficulty is the high cost and technical complexity associated with single-cell technologies, including library preparation and the need for accurate cellular barcoding to ensure the scalability of these methods to handle large sample sizes and high-throughput screening. Additionally, there is a need for standardized protocols and benchmarks to ensure reproducibility and comparability of results across different scRNA-seq studies and platforms (16). To address this, we evaluate three different high throughput scRNA-seq methods that enable cellular barcoding to track cell-drug interactions in the context of the *ex vivo* drug screening.

## MATERIAL AND METHODS

### Cell culture

The human pre-B ALL cell line Reh (DSMZ, ACC 22) was maintained in RPMI-1640 medium (Sigma, R0883) supplemented with 2 mM L-Glutamine (Sigma, G7513), 100 U/mL Penicillin, and 100 µg/mL Streptomycin (Sigma, P0781), along with 10% heat-inactivated fetal bovine serum (HIFBS; Sigma F9665). Cells were cultured at 37°C in humidified air (95%) and 5% CO2. Cells were split twice a week, with the first two subcultures after thawing the cell line supplemented with an additional 10% HIFBS in the culture media to enhance cell growth. The Reh cell line was verified by karyotyping (https://doi.org/10.1101/2023.03.08.531483) and the Eurofins human STR profiling cell line authentication service.

### *Ex vivo* drug treatment

The Fluorometric Microculture Cytotoxicity Assay (FMCA) was conducted following previously established protocols (9, 17–19). Briefly, Reh cells were resuspended in cell culture media and seeded into a 384-well plate at a density of 7000 cells in 45 µL of media per well (0.156 x 10^6^ cells/mL). A total of 41 compounds, including positive and negative controls, were added to the plate using an Echo 550 liquid handler (Beckman Coulter, Brea, CA, USA), following a 1:3 dilution series in a five-step gradient. Subsequently, the plates were incubated for 72 hours in a controlled environment (37 ℃, 95% humidity, 5% CO2). After the incubation period, the cellular response was assessed using a fully automated SCARA system (Beckman Coulter, Brea, CA, USA). The survival index (SI%) was determined by calculating the fraction of surviving cells relative to untreated cells. A comprehensive list of the compounds and their concentrations can be found in Supplementary Figure S1.

### FMCA prior to scRNA-seq

Reh cells were seeded in a six-well plate at a density of 0.16 × 10^6^ cells per well, with 3 mL of cell culture media added to each well. Two six-well plates were prepared during the cell culture split, with each plate containing five wells for drug treatment and one well serving as a vehicle control treated with dimethyl sulfoxide (DMSO). The selected drugs and concentrations used in the experiment were as follows: 5 µM 5-azacytidine (A2385, Sigma), 10 µM dexamethasone (D1756, Sigma), 10 µM imatinib (I-5508, LC Lab), 1 µM XRP44X (X3129, Sigma-Aldrich), and 0.56 µM fludarabine (S1391, Selleckchem). The control well was treated with 0.9% (v/v) DMSO (D5879, Honeywell). Following a 72-hour incubation (37 ℃, 95% humidity, 5% CO2), the cells were harvested for analysis.

### Cell harvesting

The FMCA and cell harvesting experiments were conducted in a consistent manner for each of the three scRNA-seq methods investigated. Following the treatment period, cells were harvested by transferring to 15 ml polypropylene tubes containing phosphate buffered saline (PBS). Prior to cell harvesting, the PBS was pre-warmed to 37 °C. The harvested cells were centrifuged at 200 g for 10 minutes, and then resuspended in 2.0 ml of PBS. The concentration and viability of cells in each treatment condition were assessed using AO-DAPI staining and analyzed on a NucleoCounter-3000 (Chemometec).

### Parse Biosciences Evercode library preparation (Parse)

The cells recovered from each treatment condition (range 616k-1,720k) were fixed according to the manufacturer’s protocol (Parse Biosciences) including 0.5% BSA in the fixation solution. After the fixation the cell concentration was measured for each suspension before being stored at −80°C. For barcoding, cell suspensions with equal number of cells from each treatment condition were thawed and split to two wells of the first barcoding plate of an Evercode whole transcriptome V1 mini kit (Parse Biosciences). Barcoding and sequencing library generation were performed according to the manufacturer’s protocol, generating two sub-libraries, with a target of 5,000 cells each (10,000 cells in total). Prior to library generation the concentration of the pooled and barcoded cell suspension was evaluated by trypan blue (EVE cell counter) and propidium iodide label (Cellometer K2). The input cell concentration of the first sub-library (P1) was based on the trypan blue estimate and the second sub-library (P2) was based on the propidium iodide label estimate. The two final sequencing sub-libraries were evaluated on a TapeStation (Agilent) and quantitative real time PCR (qPCR) before being pooled together at equimolar concentrations.

### MULTI-seq library preparation (MULTI-seq)

A volume equivalent to 500k cells from each treatment condition was transferred to six 15 ml tubes, and PBS was added to achieve a total volume of 5 ml. The cells were isolated by centrifugation at 200 g for 10 min and subsequently transferred to loBind 1.5 ml Eppendorf tubes. Barcode labeling was performed using lipid modified oligos (LMO, Sigma) as previously described (20). The processing was carried out in six parallel tubes, optimizing wash steps and volumes to minimize handling time and cell loss. The final resuspension of cells was performed in PBS supplemented with 1% BSA and 0.5 U/µl RNase inhibitor, using a low volume of 100 µl/sample. The LMO-barcoded samples were subsequently pooled equally by volume. Dead cells were removed by incubating with DAPI for 5 minutes, followed by negative selection using gentle sorting (Miltenyi MACS Quant Tyto). The concentration and viability of the purified cell suspension pool were evaluated using AO-DAPI staining (NucleoCounter-3000). The suspension pool with a volume equivalent to a target of 6,000 cells was immediately processed on one lane of a Chromium next GEM Single cell 3’ gene expression v3.1 dual index kit (10x Genomics) following the manufacturer’s protocol. During the initial cDNA purification step, the fraction containing the short cDNA, which contained the barcode libraries, was used for generating a barcode library according to the MULTI-seq protocol. The gene expression library and barcode library were evaluated using a TapeStation (Agilent) and qPCR before being pooled.

### 10x Genomics Single Cell Gene Expression Flex library preparation (10x Fixed)

Cells from each treatment condition (518k–1390k cells) were fixed with formaldehyde and additive according to the Chromium Next GEM Single Cell Fixed RNA Sample Preparation Kit (10x Genomics). Fixation was performed at +4°C overnight. After buffer exchange, the fixed cells were stored at −80°C until processing. A four barcode (BC) version of Chromium Fixed RNA Kit, Human Transcriptome (10x Genomics) was used for subsequent barcoding of the six different samples, i.e., three BC sets were combined in each 10x reaction (n = 2 libraries). The post hybridization washes were done individually for each barcoded sample, according to the manufacturer’s protocol. Cell counting was performed using propidium iodide labeling (Cellometer K2). A target of 2,500 cells per treatment condition were combined for a total of 7,500 cells per library using a Chromium iX instrument. The two resulting libraries were generated according to the manufacturer’s protocol. The libraries were evaluated using a TapeStation (Agilent) and qPCR before being pooled.

### Sequencing

Sequencing was performed at the SciLifeLab National Genomics Infrastructure in Uppsala, Sweden. Each pool of libraries was sequenced independently by method on one lane of a NovaSeq6000 S Prime (SP) flowcell (Illumina), for three SP lanes in total. A paired-end read setup of 100bp-10bp-10bp-100bp (Read1-index1-index2-read2) for Parse and MULTI-seq or 150bp-10bp-10bp-150bp for 10x Fixed, was used with v1.5 sequencing chemistry. Base calling was performed with RTA v3.3.4 and bcl2fastq 2.20.0.422.

### Data processing

The data from chromium-based methods (MULTI-seq and 10x Fixed) were demultiplexed and mapped with Cellranger v7.0.1 using a GRCh38 reference provided by 10x Genomics (refdata-gex-GRCh38-2020-A). For MULTI-seq, the gene expression analysis was run in parallel with feature barcode analysis. The feature barcode matrix was subsequently loaded into R where the two assays were merged and samples were demultiplexed using HTODemux from the Seurat package (21). Cells classified as negatives or doublets were removed. Parse data was demultiplexed and mapped with the ParseBiosciences-Pipeline v0.9.6p using the GRCh38 reference (refdata-gex-GRCh38-2020-A). The unfiltered count matrix was used for downstream analysis. Further analysis of the raw count matrices was performed in R using Seurat, harmony and ggplot2 packages in R. The inflection point of the total count versus barcode rank curve was determined for each library using the barcodeRanks function from DropletUtils (22). Cells with a total UMI count lower than the inflection point and cells with >10% of the total counts originating from mitochondrial genes were removed (Supplementary Figure S2). Following cell filtering, libraries from the Parse and 10x Fixed experiments were each merged to create a single dataset per method. Log-normalization and scaling was performed with Seurat v4 and variance stabilizing transformation was performed with SCTransform v2 without covariates (23, 24). The data from the three scRNA-seq methods were combined using Seurat prior to clustering and visualization.

### Data analysis

#### Down sampling

The libraries were downsampled using the downsampleReads function from DropletUtils (22). Down-sampling was performed on a cell-by-cell basis generating a downsampled datasets where all cells with read counts above the down sampling target had the same total read count and cells with read counts below the down sampling target were discarded. The downsampleReads function uses the hdf5 molecule information file generated by the Cellranger pipeline as input for MULTI-seq and 10x Fixed. For the Parse libraries, the tscp assignment.csv.gz was converted to hdf5 format with a custom a python script.

#### Dropout rate

Dropout was calculated in a similar approach to Ziegenhain et al (25): First, we randomly subsampled 500 DMSO-treated cells per scRNA-seq method for a total of 1,500 cells. Next, we selected the genes detected (non-zero count) in at least 25% of all cells by any scRNA-seq method. Finally, we calculated the dropout rate by scRNA method and by gene as the proportion of cells with zero counts.

#### Cell cycle scores

Cell cycle scores were computed using the CellCycleScoring function and marker genes classify cells into G1, G2M or S states in Seurat (21).

#### Differential gene expression

Differential gene expression was determined using MAST (26). Differentially expressed genes were determined by comparing each drug treatment separately to control (DMSO).

#### Pathway analyses

The R-package clusterProfiler was used for over-representation analysis for GO biological process, WikiPathways, KEGG and CMAP on genes determined to be differentially expressed (fold change >0.75, adjusted p<0.01) as compared to DMSO controls (27, 28). Significant overlaps (FDR q-values < 0.05) between differentially expressed genes and Hallmark gene sets were computed for each sequencing method by Gene Set Enrichment Analysis (29).

### Cell cycle analysis by DNA stain

Reh cells were seeded in a six-well plate at a density of 0.16 × 10^6^ cells per well, with 3 mL of cell culture media added to each well and treated with 5 µM 5-azacytidine (A2385, Sigma), 10 µM dexamethasone (D1756, Sigma), 10 µM imatinib (I-5508, LC Lab), 0.56 µM fludarabine (S1391, Selleckchem) or 0.9% (v/v) DMSO (D5879, Honeywell). Following a 72-hour incubation (37 ℃, 95% humidity, 5% CO2), the cells were harvested (see Cell Harvesting above). For each treatment 0.5 x 10^6^ cells were lysed and stained with 10 μg/ml DAPI according to the “Two-step cell cycle analysis” application note (Chemometec). The fluorescence distribution was quantified in a NC-3000 (Chemometec) image cytometer.

## RESULTS

### Functional drug resistance testing by FMCA

We performed a systematic methods comparison study of three approaches for multiplexing scRNA-seq experiments in an *ex vivo* drug screening model system (FMCA). To identify drugs suitable for the method comparison, we initially assessed the *ex vivo* cellular drug resistance of the glucocorticoid-resistant E/R+ ALL Reh cell line using FMCA. A panel of 41 drugs from diverse classes, including targeted drugs, glucocorticoids, and chemotherapeutic agents, was employed at varying concentrations (Supplementary Figure S1). Based on the dose-response curves after 72 hours of treatment, five drugs were selected for further investigation of their transcriptional effects (Figure 1). The chosen drugs were as follows: 5-azacytidine, a nucleoside analogue that inhibits DNA methyltransferase, resulting in DNA hypomethylation and cell death at high doses (30, 31). Dexamethasone, a glucocorticoid, was included as a negative control since the Reh cell line is known to exhibit glucocorticoid resistance (32, 33). Imatinib, a tyrosine kinase inhibitor used in the treatment of ALL patients with ABL-class fusions, although the Reh cell line does not possess such fusions; its inclusion was based on the responsive profile (Figure 1). XRP44X, an inhibitor of Ras-activated Elk3 transcription-factor activity, was selected due to a previous observation in t(12;21) E/R ALL (34). Fludarabine, a purine analog, is currently employed in the treatment of high-risk ALL patients as part of conditioning before stem cell transplantation and lymphodepletion before CAR-T therapy (35). For each drug, a concentration was determined for subsequent single-cell analysis, aiming to achieve a SI% of ∼50% to balance between inducing a cellular response, while maintaining a sufficient proportion of surviving cells for robust assessment of transcriptional effects.

**Figure 1.**
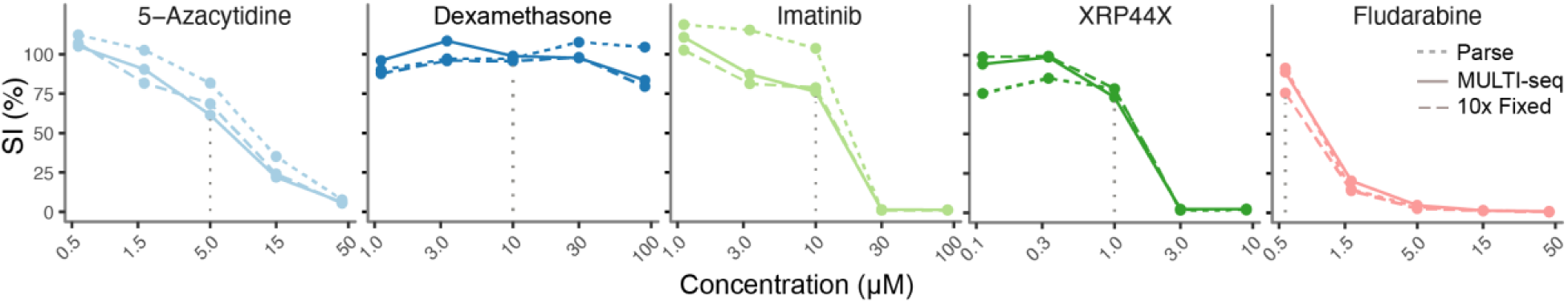
Fluorometric Microculture Cytotoxicity Assay (FMCA) survival indexes (SI%) for five selected drugs. The SI% (y-axis) after exposed to 72 hours treatment at a range of concentrations (x-axis). Each dot represents the mean SI% measured from two replicates. The dotted vertical line denotes the concentration selected for single-cell transcriptomic analyses (scRNA_seq). The results from the FMCA experiment used to generate each of the scRNA-seq datasets are indicated according to the legend in the right side of the panel.

### Multiplexed scRNA-seq library preparation

We evaluated the *ex vivo* transcription profiles of the Reh cells after treatment with the five aforementioned drugs and DMSO controls. Three library preparation protocols were tested: 1) the Parse Biosciences Evercode Whole Transcriptome mini v1 (Parse) (36), 2) the cellular LMO-barcoding approach in combination with scRNA-seq using the 10x Genomics Chromium 3’ GEX V3 protocol (MULTI-seq) (20), and 3) the Chromium Single Cell Gene Expression Flex kit from 10x Genomics (10x Fixed).

To capture subtle differences in cell states by each drug condition, we aimed to profile 6,000–15,000 cells with each scRNA-seq protocol (1,000 - 2,500 cells per drug condition), based on each kit’s ability to incorporate a drug-specific cellular barcode to mark surviving cells according to which drug they were exposed. Due to protocol differences and restraints in number of sample barcodes available for each kit, the library set-up for each method was slightly different. For Parse we split each sample into two initial barcoding wells (technical replicates) and in the final step before library amplification the pooled cells were split into two sublibraries containing all six conditions (two sub-libraries: P-1 and P-2). For MULTI-seq we performed a single experiment (M-1). For 10x Fixed we used three barcodes (corresponding to 5-azacytidine, XRP44X, dexamethasone) in library F-1 and three barcodes (corresponding to imatinib, fludarabine and DMSO control) in library F-2. Differences in sequencing depth, UMIs and number of genes detected prior to cell filtering and library merging were observed between the Parse P-1 and P-2 sublibraries as well as the 10x Fixed F-1 and F-2 libraries (Supplementary Figure S2-3). The differences in sequencing depth per cell of P1 and P2 sublibraries was correlated with cell concentration measurement as described below. After removing non-cell UMIs, cells with >10% mitochondrial reads, and merging the sub-libraries we characterized 27,327 cells in total: 7,225 cells with the Parse method, 10,516 with MULTI-seq, and 9,586 with 10x Fixed (Table 1, Supplementary Figure S3).

**Table 1.** Single-cell RNA-sequencing libraries and statistics.

| Method | Parse |  | MULTI-seq | 10x Fixed |  |
| --- | --- | --- | --- | --- | --- |
| Sub-library name | P-1 | P-2 | M-1 | F-1 | F-2 |
| Number single cells targeted | 5 000 | 5 000 | 6 000 | 7 500 | 7 500 |
| Number single cells identified | 2 296 | 4 929 | 10 516 | 4 257 | 5 329 |
| Total reads sequenced / cells identified | 111 k | 42 k | 43 k | 41 k | 62 k |
| Median reads per cell | 26 771 | 9 605 | 27 483 | 32 241 | 49 279 |
| Median genes per cell | 3 772 | 2 698 | 4 167 | 6 262 | 6 927 |

### Evaluation of library complexity

Since several approaches were used for cell counting prior to library preparation, we estimated how well each method performed. In our hands, the propidium iodide labeling used to count cells in P-2, F-1 and F-2 more accurately estimated the cell concentration, while quantification by trypan blue overestimated the cell concentration, resulting in underloading of library P-1, while AO-DAPI underestimated the cells loaded in M-1, resulting in overloading. We therefore conclude that accurate cell counting is important for downstream data quality and propidium iodide labeling was the most reliable method for determining cell concentration prior to scRNA-seq library preparation for fixed cells.

The proportion of raw reads retained for downstream analysis after demultiplexing and quality filtered varied by scRNA-seq method (Figure 2a). The most sequencing reads were retained in 10x Fixed (85.5 %), followed by MULTI-seq (60.6 %) and Parse (51.6 %). We further examined read retainment through the different steps of the pipeline For the Parse and MULTI-seq libraries, the majority of reads (>25%) were removed during demultiplexing, due to incomplete or non-matching sample barcode sequences (Supplementary Figure S4). The proportion of filtered reads that mapped to exonic, intronic or intergenic regions also varied between the libraries generated by the three methods, which likely reflects differences in the method design (Figure 2b). The 10x Fixed approach resulted the highest proportion of exonic reads (98.7 %), consistent with the assay using probes targeting exons. MULTI-seq, based on the 10x Genomics 3’ assay that uses oligo (dT) primers, produced 75% reads mapped to exons in line with what others have shown previously (16). Parse uses random primers in addition to oligo(dT), resulting in the majority of reads mapping to intronic regions (60.4 %), which is in line with what other reports using this method have observed (https://doi.org/10.1101/2023.06.28.546827).

**Figure 2.**
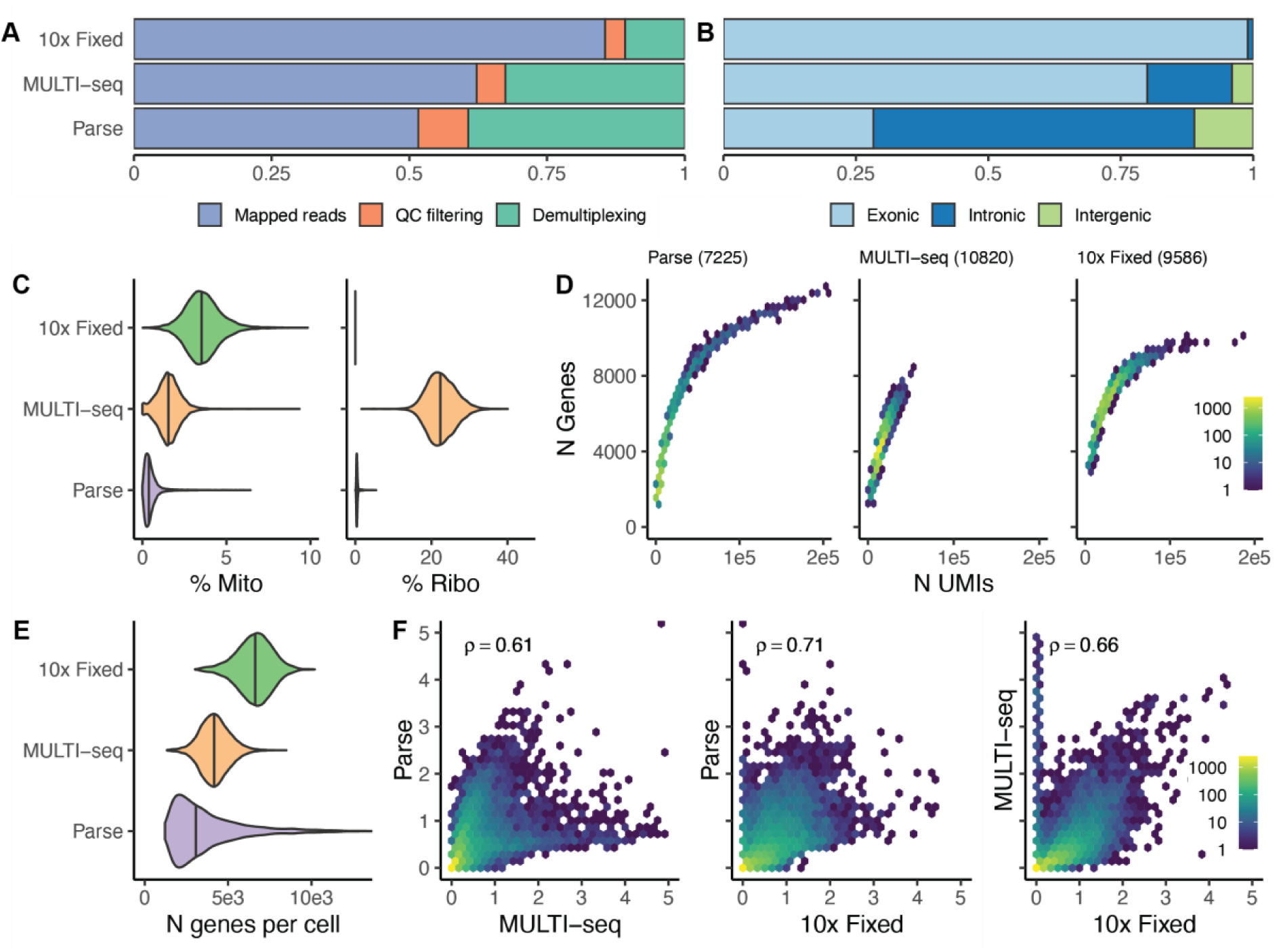
Quality metrics of single cell RNA sequencing data generated from three different library preparation methods (Parse, MULTI-seq, 10x Fixed). (A) Stacked bar plot showing the percentage of sequencing reads lost across each step of data processing. (B) Proportion of mapped reads that map to exonic, intronic and intergenic regions by scRNA-seq method. (C) Violin plots of the of mitochondrial (left) and ribosomal (right) read fractions separated by scRNA-seq method. (D) The number of genes detected (y-axis) plotted against the number of UMIs (x-axis) for each scRNA-seq library preparation method. The numbers are visualized as hexagonal binning plots, the color indicates the number of cells contributing to each hexagonal bin according to the figure legend scale to the right of the panel. The total number of cells captured by each method is indicated in brackets. (E) Violin plot showing the distribution of genes detected per cell for each scRNA-seq method. (F) Pairwise scatterplot of the pseudobulk gene expression level per gene and method. The Pearson’s correlation coefficient (ρ) is indicated in each panel.

The presence of a high mitochondrial read fraction in scRNA-seq data can indicate cellular stress or cell death, while the ribosomal read fraction can vary across cell state and may be an indication of RNA degradation (37, 38). To assess these parameters, we compared the proportion of reads mapping to mitochondrial and ribosomal genes across all cells by scRNA-seq method. Notably, the Parse method exhibited the lowest mitochondrial (median 0.39% Figure 2c) and ribosomal (median = 0.42%) read fraction in the filtered data as well as prior to the initial UMI filtering step (Supplementary Figure S3). This may be attributed to the multiple washing steps incorporated in the Parse workflow, which likely minimizes ambient RNA and the impact of mitochondrial/ribosomal gene expression. The MULTI-seq library contained a low mitochondrial read fraction (median 1.55%), but higher ribosomal read fractions (median 22.37%). The low fraction of mitochondrial reads likely is a result of the dead cell removal step used for the MULTI-seq library, where the cells were >99% viable prior to cell capture. The Chromium 3’ chemistry, used in MULTI-seq captures a higher proportion of ribosomal gene reads compared to other scRNA-seq methods (https://doi.org/10.1101/2023.06.28.546827)(39). The 10x Fixed libraries contained the highest mitochondrial read fraction (median 3.53%), even though the method had the shortest time between cell isolation and fixation. Unlike the other protocols, the10x Fixed pre-processing steps did not include dead cell removal nor extensive cell washes, which may attribute to this difference. No reads mapping to ribosomal genes were detected in the10x Fixed libraries as the there are no probes for ribosomal genes included in the design. Despite our experimental design involving the use of five drug treatments aimed at inducing cell death, we did not observe an increased ratio of mitochondrial genes when comparing drug-treated cells to control (DMSO) cells within each method (Supplementary Figure S5).

The relationship between the number of UMIs sequenced and genes detected per cell differed by method (Figure 2d), with Parse achieving the cells with the greatest total number of genes detected (maximum 15,594 genes in a single cell). However, taking the average number of genes detected per cell across the all the sequenced cells as a measure of sensitivity, the 10x Fixed approach achieved the highest sensitivity (mean±SD = 6,572±1,064 genes/cell), followed by MULTI-seq (mean±SD = 4,199±881), while Parse showed the lowest sensitivity in detecting genes, as well as greatest variability between the cells (mean±SD = 3,690±2,140) (Figure 2e). To further examine the agreement between methods, we performed pairwise comparisons of the average expression of each gene by method for the DMSO control cells (Figure 2f). Overall, moderate correlations were observed between the methods across all of the treatments (ρ range = 0.61 −0.74), with Parse and 10x Fixed achieving the highest concordances (Supplementary Table S1).

### Down sampling

To fairly compare the efficiency of mRNA capture between protocols, we downsampled the sequencing reads per cell to a common depth and stepwise-reduced fractions (80,000-2,500 reads per cell) for each of the three scRNA-seq methods, revealing large differences in sensitivity (Figure 3a-b). The 10x Fixed approach consistently detected more genes per cell than the other methods, especially at lower sequencing depths. Once the sequencing depth exceeded 60,000 reads per cell the number of genes detected per cell was similar between the 10x Fixed and Parse libraries, albeit in a significantly smaller proportion of cells in the Parse libraries. This was further visualized in UMAP plots where data the original dataset was compared to data downsampled to 20,000 and 5,000 reads per cell. The MULTI-seq and 10x Fixed approaches displayed a more even distribution of genes detected per cell and the 10x Fixed approach detected the most genes across the cell population (Figure 3c-e). To further compare between the libraries, we sub-sampled the DMSO cells from the three scRNA-seq datasets (20k and 5k reads/cell) to equal number of cells per library to avoid bias in differing number of cells and subsequently calculated the average expression, variance, and dropout rate for the geneset present in 25% of the cell subset (Figure 3f). In this comparison, 10x Fixed displayed highest average expression and standard deviation, as well as lowest dropout rates across the downsampled datasets (27.0 – 65.4%), while MULTI-seq had the lowest overall expression levels as well as the highest median dropout rates (53.0 – 80.4%). In summary, 10x Fixed detects the common sets of genes in more cells than the other two methods.

**Figure 3.**
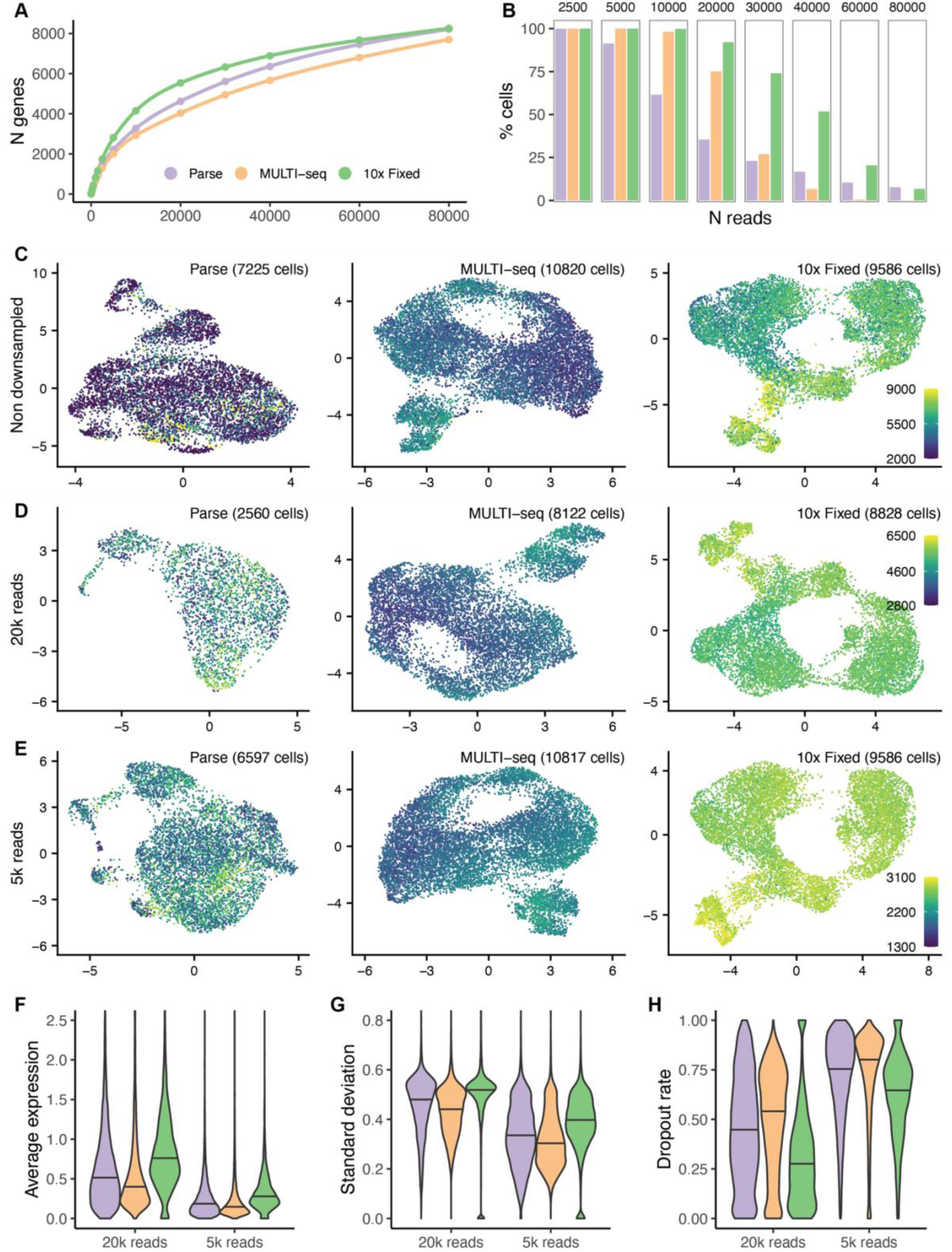
Gene detection and drop out in downsampled data. (A) The median number of genes detected per cell (y-axis) by downsampled sequencing depth in reads per cell (x-axis). The single-cell RNA-sequencing (scRNA-seq) library preparation method is indicated by color according the legend. (B) The proportion of cells with sufficient sequencing reads to be compared at each downsampled read depth, color-coded by scRNA-seq method. (C-E) UMAPs by scRNA-seq. Each cell is color coded by the number of genes detected in each cell, according to the legend on the right side of each panel. The number of cells in each UMAP panel is indicated in the brackets. The UMAPs in panel C are generated from non-downsampled data. The UMAPs in panel D are generated from downsampled data to 20,000 reads per cell. The UMAPs in panel E are generated from downsampled data to 5,000 reads per cell. (F-G) The average expression (F) and standard deviation (G) calculated using across 7,766 genes detected in 25% of 1,500 sub-sampled DMSO-treated cells (500 cells per scRNA-seq method) downsampled to 20k reads/cell and the 3,627 genes detected in the same dataset, downsampled to 5k reads per cell. (H) The dropout rate is calculated by method and gene as the proportion of the 500 cells in which the gene is not detected (zero counts).

### Drug-specific transcriptional profiles

Next, we examined the influence of specific drug conditions on gene expression profiles of the single cells recovered by the three methods. Unsupervised clustering of co-normalized data revealed that these sequencing approaches captured orthogonal aspects of the data in a highly complementary way (Figure 4a). The cells exposed to the drugs revealed a largely overlapping pattern with DMSO treated control cells, except for the cells exposed to fludarabine, where the majority of the fludarabine treated cells displayed a different transcriptional state and were confined to one cluster (Figure 4b). This pattern was evident in each of the three scRNA-seq methods (Supplementary Figures S6-8). Each cell was further assigned into G1, S or G2M phases of the cell cycle based on cell cycle scoring in Seurat (21) (Figure 4c, Supplementary Figures S6-8), which revealed that the majority of fludarabine treated cells displayed a transcriptional profile related to the S-phase of the cell cycle (Figure 4d). We examined the cell cycle with orthogonal analysis with DNA stain intensity, which confirmed that a large proportion of the cells appear to be in S-phase, with few cells in G2/M, further supporting that the fludarabine treated cells were in cell cycle arrest (Supplementary Figure S9).

**Figure 4.**
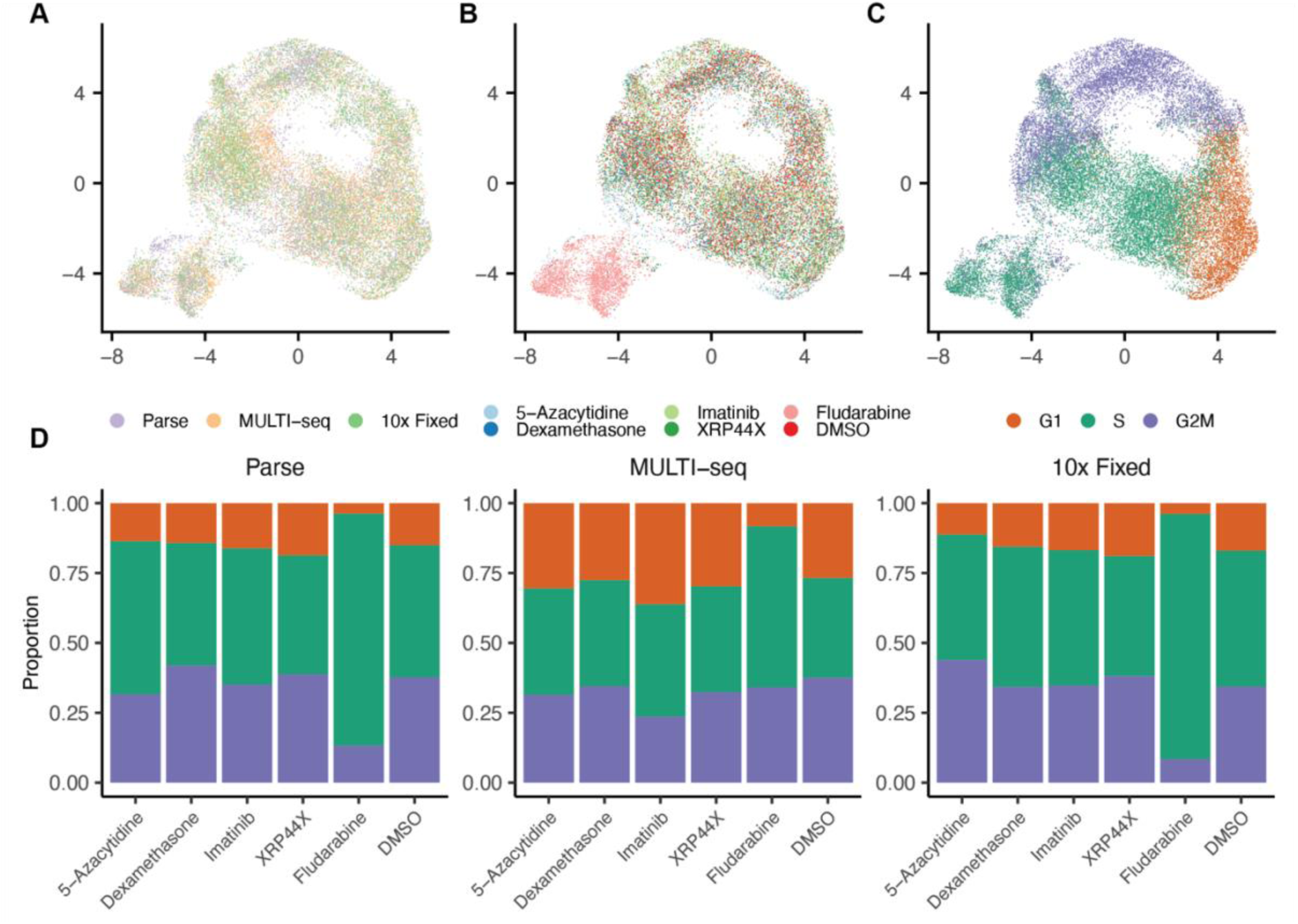
Global gene expression in the dataset. (A-C) UMAPs of the merged dataset of 27,631 cells from the Parse, MULTI-seq and 10x Fixed libraries. Each dot in the graph represents a single cell. The UMAPs are color coded by library preparation method (A), drug condition (B), and by cell cycle phase marker gene expression (C). The color key is indicated at the bottom of each respective panel. (D) Stacked barplots indicating the proportions of cells annotated to G1-, S- or G2/M-phases of the cell cycle by scRNA-seq method.

### Ability of scRNA-seq methods to detect drug-specific differential gene expression

To further examine the underlying transcriptional profiles and how they differed between methods and treatment conditions, differential gene expression analyses were performed using MAST (26), by scRNA-seq method, comparing cells between drug-treated condition and DMSO controls. We used a stringent cut-off (absolute log fold change >0.75, adjusted p<0.01) for determining differentially expressed genes (DEGs). The resulting DEGs of were compared for each of the treatments (Figure 5a). As expected due to the glucocorticoid resistance of the Reh cell line (33), few or no DEGs were identified in the dexamethasone treated cells (Supplementary Table S2). A small number of DEGs were detected as differentially expressed in the 5-azacytadine, Imatinib, and XRP44X treated cells (Supplementary Table S3-S5), and the largest number of DEGs were found in the fludarabine treated cells: 409 by 10x Fixed, 70 by MULTI-seq, and 27 by Parse (Supplementary Table S6). Gene Set Enrichment Analysis (29) of fludarabine-treated cells suggested a high concordance of Hallmark gene sets enriched in differentially expressed genes called by any of the three scRNA-seq methods (Table 2, Figure 5b). In particular, upregulation of p53 pathway signaling members (e.g. *CDKN1A*, *TRIB3*, *BTG2*, *DDIT3/4,* and *FDXR*; FDR q-value = Parse 4.17 x 10^-35^; MULTI-seq 3.49 x 10^-35^; 10x Fixed 7.12×10^-33^) and downregulation of MYC targets (e.g. *MYC*, *HSPD1*, *NPM1*, *PRMT3*, *SRSF1/2*; FDR q-value = 2.37×10^-22^; 2.09×10^-22^; 2.43×10^-22^) were among the top enriched gene sets present in treated cells. Among the genes involved in these gene sets, we particularly detected p53-dependent mediators of apoptosis, as well as genes involved in DNA damage response and oxidative stress. We additionally observed a significant downregulation of genes involved in G2/M-mediated cell cycle progression (FDR q-value = 1.56×10^-8^; 1.47×10^-8^; 8.73×10^-10^), supportive of our previous functional observations where fludarabine treatment induces cell cycle arrest in Reh cells. To further characterize the interaction between genes dysregulated by fludarabine treatment, we performed a STRING network analysis (40) on the set of significant differentially expressed genes (Supplementary Figure S10). Here, k-means clustering indicated strong interactions between six clusters of encoded proteins, including p53-mediated regulators of apoptosis, DNA damage response, and oxidative stress. Altogether, these results support the ability of the tested scRNA-seq methods to detect biologically relevant pathways associated with drug perturbations in ALL.

**Figure 5.**
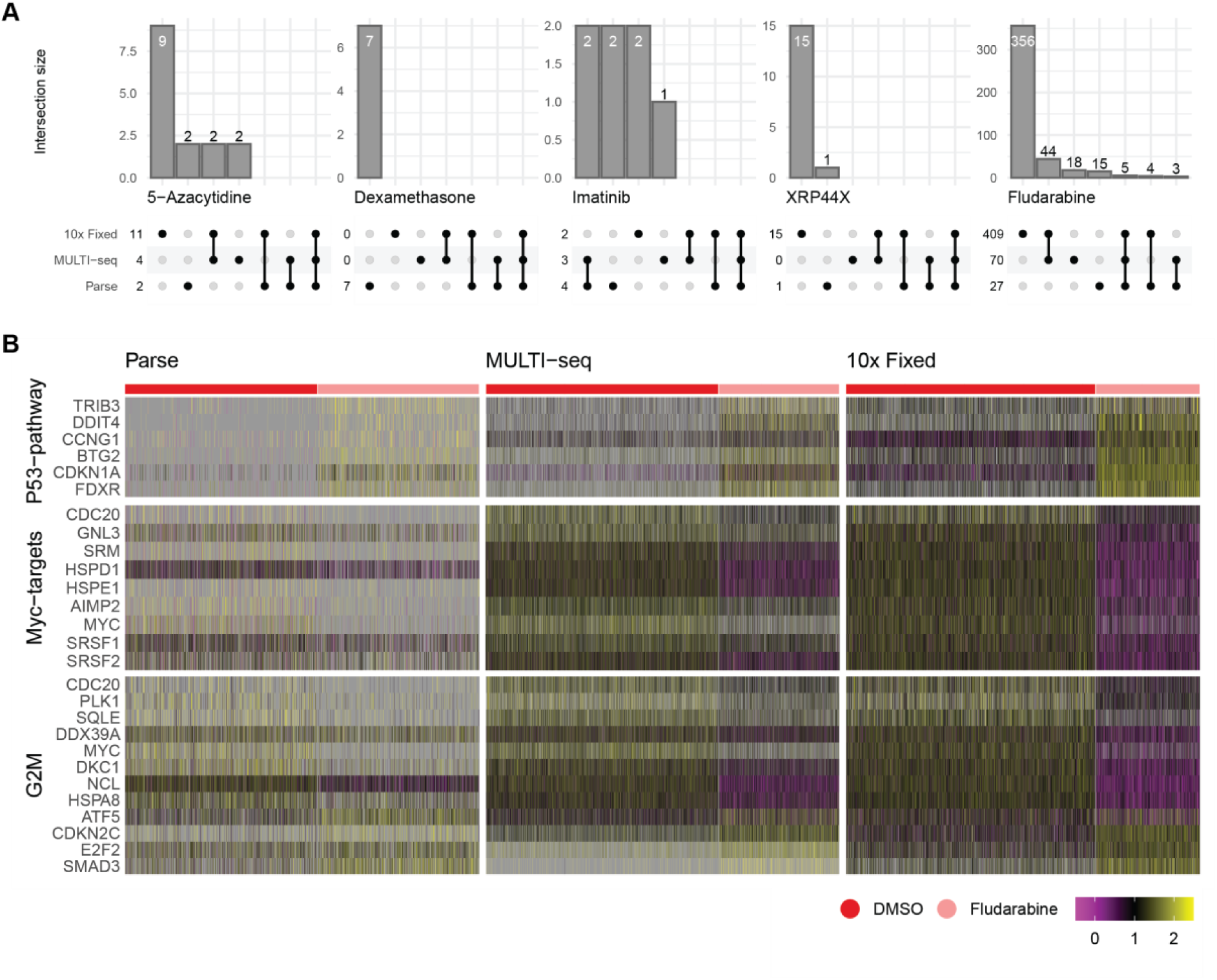
Differential gene expression analyses. (A) Upset plots show the number of genes with significant (p<0.01) altered expression (absolute log fold change >0.75) in the cells surviving each of the five treatment conditions compared against the DMSO-treated control cells. To the left of each plot the numbers of significantly differentially expressed genes are indicated and the upper bars indicate the numbers of genes common (or unique) between the methods. (B) Heatmap of differentially expressed genes in the enriched pathways in the fludarabine-treated cells. The cells are ordered along the x-axis by scRNA-seq method and then by control (DMSO, red) and fludarabine treated (pink). The differential expressed genes are ordered along the y-axis and grouped according to the pathway. The color key for the gene expression levels is indicated at the bottom right side of the panel. Purple indicates low expression, yellow high expression, and grey is missing data.

**Table 2.** Gene set enrichment analysis.

| Hallmark Gene Set Name | Description | # Genes in Gene Set (K) | 10x Fixed |  |  |  | MULTI-seq |  |  |  | Parse |  |  |  |
| --- | --- | --- | --- | --- | --- | --- | --- | --- | --- | --- | --- | --- | --- | --- |
|  |  |  | # Genes in Overlap (k) | k/K | p-value | FDR q-value | # Genes in Overlap (k) | k/K | p-value | FDR q-value | # Genes in Overlap (k) | k/K | p-value | FDR q-value |
| P53 Pathway | Genes involved in p53 pathways and networks. | 200 | 38 | 0.19 | 8.34E-37 | 4.17E-35 | 38 | 0.19 | 6.98E-37 | 3.49E-35 | 34 | 0.17 | 1.42E-34 | 7.12E-33 |
| MTORC1 signaling | Genes up-regulated through activation of mTORC1 complex. | 200 | 28 | 0.14 | 1.42E-23 | 2.37E-22 | 28 | 0.14 | 1.25E-23 | 2.09E-22 | 26 | 0.13 | 1.46E-23 | 2.43E-22 |
| MYC Targets V1 | A subgroup of genes regulated by MYC - version 1 (v1). | 200 | 28 | 0.14 | 1.42E-23 | 2.37E-22 | 29 | 0.145 | 7.08E-25 | 1.77E-23 | 26 | 0.13 | 1.46E-23 | 2.43E-22 |
| MYC Targets V2 | A subgroup of genes regulated by MYC - version 2 (v2). | 58 | 15 | 0.2586 | 1.87E-17 | 2.34E-16 | 16 | 0.2759 | 4.60E-19 | 5.75E-18 | 14 | 0.2414 | 3.05E-17 | 3.81E-16 |
| Cholesterol homeostasis | Genes involved in cholesterol homeostasis. | 74 | 15 | 0.2027 | 9.90E-16 | 9.90E-15 | 15 | 0.2027 | 9.24E-16 | 9.24E-15 | 12 | 0.1622 | 9.46E-13 | 7.88E-12 |
| E2F Targets | Genes encoding cell cycle related targets of E2F transcription factors. | 200 | 21 | 0.105 | 2.07E-15 | 1.73E-14 | 21 | 0.105 | 1.89E-15 | 1.57E-14 | 20 | 0.1 | 3.83E-16 | 3.83E-15 |
| TNFA signaling via NFKB | Genes regulated by NF-kB in response to TNF [GeneID=7124]. | 200 | 17 | 0.085 | 3.01E-11 | 2.15E-10 | 17 | 0.085 | 2.80E-11 | 2.00E-10 | 11 | 0.055 | 8.54E-07 | 4.98E-06 |
| G2M Checkpoint | Genes involved in the G2/M checkpoint, as in progression through the cell division cycle. | 200 | 15 | 0.075 | 2.50E-09 | 1.56E-08 | 15 | 0.075 | 2.34E-09 | 1.47E-08 | 15 | 0.075 | 1.22E-10 | 8.73E-10 |
| Inflammatory response | Genes defining inflammatory response. | 200 | 14 | 0.07 | 2.05E-08 | 1.14E-07 | 14 | 0.07 | 1.93E-08 | 9.65E-08 | NA | NA | NA | NA |
| Apoptosis | Genes mediating programmed cell death (apoptosis) by activation of caspases. | 161 | 12 | 0.0745 | 1.06E-07 | 5.29E-07 | NA | NA | NA | NA | 10 | 0.0621 | 8.96E-07 | 4.98E-06 |
| Allograft rejection | Genes up-regulated during transplant rejection. | 200 | NA | NA | NA | NA | 14 | 0.07 | 1.93E-08 | 9.65E-08 | 10 | 0.05 | 6.28E-06 | 2.85E-05 |

## DISCUSSION

In this study, we conducted a comprehensive assessment of three multiplexed experimental approaches to investigate transcriptional changes in individual cells during cytotoxic *ex vivo* drug screening (FMCA). Our aim was to compare the performance of three single-cell scRNA-seq methods using reference leukemia (Reh) cells treated with various cytotoxic drugs. By evaluating a wide range of performance metrics, we were able to identify technical strengths and limitations intrinsic to each approach. Our findings demonstrate, through the example of fludarabine-treated cells, that any of the scRNA-seq methodologies could robustly discern mechanistic underpinnings of the drug in ALL cells.

Each of the three scRNA-seq methods employed in our study generated high-quality data for single-cell gene expression profiling. The Parse and 10x Fixed methods offered the advantage of cell fixation, allowing for potential sample storage and collection at multiple time points. This feature may be particularly beneficial for studies involving delicate or easily disrupted cell types (41). On the other hand, the MULTI-seq protocol has the greatest potential for achieving ultra-high multiplexing (20) and thus lower costs, although it required a larger number of input cells and was more time-consuming compared to the other methods. It is worth noting that minimizing the inclusion of ambient RNA from dying cells during multiplexing is crucial for obtaining optimal results when using LMO or similar antibody-based cell hashing protocols, as suggested previously (42, 43). In our hands, the additional washing steps and the need for performing a dead cell removal step after labeling with LMOs contributed to the longer processing time until mRNA capture, which made the MULTI-seq approach the most laborious of the three methods tested. At present, the 10x Fixed protocol is most limited in terms of number of barcodes available (n=16), however it is feasible to process several pooled reactions in parallel. Future endeavors focused on i) increasing the multiplexing capabilities of scRNA-seq experiments, ii) streamlining the laboratory procedure, and iii) efficiently fixing cells, will be needed to bring down the cost of fully implementing single-cell read outs into high-throughput *ex vivo* drug screening.

In terms of sequencing efficiency and gene detection sensitivity the 10x Fixed approach outperformed the other two methods. A higher number of genes per cell (mean = 6.5k) were consistently detected with 10x Fixed, which is in line with a previous study utilizing the same method (https://doi.org/10.1101/2023.04.25.538273). Our differential expression analysis further underscored findings revealing more differential expressed genes using the 10x Fixed approach. Despite that the Parse approach achieved the cells with the highest number of UMIs sequenced and genes detected, this was observed in only a small fraction of individual cells.

Our study revealed that treatment with fludarabine elicited the most pronounced transcriptional changes, exhibiting a large number of differentially expressed genes that converge on p53 pathway signaling and Myc transcription factor activity. Fludarabine treatment is known to induce a p53-dependent transcriptional responses in chronic lymphocytic leukemia (45,46), and our study highlights a similar mechanism of action in an additional hematological malignancy model. Although we observed only minimal overlap in differentially expressed genes between the scRNA-seq method, of the genes identified, *DDB2*, was consistently detected by all three methods and is known to be induced by p53. Another gene, *CDKN1A*, which encodes the protein p21, was also upregulated and is involved in cell cycle arrest (44) and potentially inhibits apoptosis (45). *CDKN1A* is highly expressed in primary leukemic cells at relapse and, when overexpressed in Reh cells, has been associated to an increased resistance to cytarabine (12), an antimetabolite similar to fludarabine. We additionally detected a consistent downregulation of Myc transcriptional targets in fludarabine-treated cells. A recent drug repurposing effort found fludarabine phosphate to suppress *MYCN* signaling in neuroendocrine prostate cancers (46). These findings motivate further exploration of MYC transcriptional activity as a biomarker of response to fludarabine and other DNA synthesis inhibitors. Unexpectedly, our study did not uncover the anticipated analogous transcriptional perturbations across the remaining drugs (34). Several factors may contribute to this observation. One possibility is that our decision to employ stringent statistical thresholds aimed at enhancing comparability between the scRNA-seq methods may have masked subtle drug-induced transcriptional responses. Additionally, the extended 72-hour incubation period may have exceeded the therapeutic window for some of these drugs, potentially leading to missed transcriptional insights. To address these potential limitations, future experiments could consider adopting time series analyses with shorter incubation durations and a more flexible approach to statistical thresholds for capturing nuanced drug-transcriptional dynamics.

While our study comprehensively assessed three multiplexed scRNA-seq methods for the purposed of *ex vivo* drug screening, there are two limitations to be discussed. In conducting this comparative analysis, we employed a conventional commercially available B-cell leukemia model (32). Although cell line biology is distinct from primary samples in many aspects, these data can potentially serve as a valuable reference and resource, facilitating future integration and comparison of results to make biological inferences by the wider scientific community (47). Primary leukemia specimens might conceivably result in higher variability, particularly in terms of increased cellular heterogeneity, mRNA expression levels, and viability, which could have added important information on the performance of the different scRNA-seq methods. However, our cell line strategy not only served to mitigate potential sources of experimental variation across the scRNA-assays, but also permitted open data sharing and dissemination. We also acknowledge the presence of other techniques that facilitate the multiplexing of cells originating from multiple conditions in a single RNA-seq library. Notable examples include transient transfection of short barcoding oligos (SBOs) (48), antibody-based cell hashing (49), the 10x Genomics 3’ Cellplex approach, among others. Consequently, our study does not cover all possible approaches in this field. Moreover, we recognize that our analyses were conducted leveraging the first version of the Parse Biosciences Evercode Whole Transcriptome method. It is plausible that the improved second iteration (V2) of the kit may yield different metrics and performance outcomes (https://doi.org/10.1101/2022.08.27.505512).

Overall, the outcomes of this study underscore both the technical disparities inherent to scRNAseq methods and the remarkable concurrence in the biological processes underpinning drug response. This level of concurrence in the scRNA-seq data is important for extending future possibilities to explore the mechanism of action for other therapeutics. In conclusion, we show that combining *ex vivo* drug screening with the high resolution of scRNA-seq is a feasible approach to uncovering mechanisms of action for drugs, bridging struggles associated with cellular heterogeneity in bulk samples, and opens an exciting opportunity for generating new biological insight in future FPM studies.

## DATA AVAILABILITY

The scRNA-seq data was deposited at the Gene Expression Omnibus (GEO, https://www.ncbi.nlm.nih.gov/geo) under accession number GSE229617.

## SUPPLEMENTARY DATA

Supplementary Data are available online.

## AUTHOR CONTRIBUTIONS

Henrik Gezelius: Conceptualization, Methodology, Investigation, Formal analysis, Validation, Visualization, Writing-original draft. Anna Pia Enblad: Conceptualization, Formal Analysis, Writing-review & editing. Anders Lundmark: Formal analysis, Data curation, Visualization. Martin Åberg: Investigation, Validation, Writing-review & editing. Kristin Blom: Methodology. Jakob Rudfeldt: Investigation, Validation. Arja Harila: Conceptualization. Amanda Raine: Conceptualization. Verónica Rendo: Formal Analysis, Writing-review & editing. Merja Heinäniemi: Writing-review & editing. Claes Andersson: Methodology, Recourses, Formal analysis, Writing-review & editing. Jessica Nordlund: Conceptualization, Recourses, Funding acquisition, Visualization, Writing-original draft.

## ACKNOWLEDGEMENTS

Sequencing was performed by the SciLifeLab National Genomics Infrastructure (NGI) SNP&SEQ unit at Uppsala University. NGI is funded by SciLifeLab, the Knut and Alice Wallenberg Foundation, and the Swedish Research Council through grant agreement no. 2019-0222. The computation analyses were enabled by resources in project [SNIC 2022/22-63] provided by the National Academic Infrastructure for Supercomputing in Sweden (NAISS) at UPPMAX, funded by the Swedish Research Council through grant agreement no. 2022-06725. We thank Katarina Tegner, Marie Söderberg, Anna Haukkala, Magnus Lindell, Elin Övernäs and Sara Ekberg from NGI for assistance with sequencing and data processing.

## REFERENCES

1. Letai, A., Bhola, P. and Welm, A.L. (2022) Functional precision oncology: Testing tumors with drugs to identify vulnerabilities and novel combinations. Cancer Cell, 40, 26–35.

2. Larsson, R., Nygren, P., Ekberg, M. and Slater, L. (1990) Chemotherapeutic drug sensitivity testing of human leukemia cells in vitro using a semiautomated fluorometric assay. Leukemia, 4, 567– 571.

3. Beaubier, N., Bontrager, M., Huether, R., Igartua, C., Lau, D., Tell, R., Bobe, A.M., Bush, S., Chang, A.L., Hoskinson, D.C., et al. (2019) Integrated genomic profiling expands clinical options for patients with cancer. Nat Biotechnol, 37, 1351–1360.

4. Fukuhara, S., Oshikawa-Kumade, Y., Kogure, Y., Shingaki, S., Kariyazono, H., Kikukawa, Y., Koya, J., Saito, Y., Tabata, M., Yoshifuji, K., et al. (2022) Feasibility and clinical utility of comprehensive genomic profiling of hematological malignancies. Cancer Science, 113, 2763–2777.

5. Ross, J.S., Wang, K., Gay, L., Otto, G.A., White, E., Iwanik, K., Palmer, G., Yelensky, R., Lipson, D.M., Chmielecki, J., et al. (2015) Comprehensive Genomic Profiling of Carcinoma of Unknown Primary Site: New Routes to Targeted Therapies. JAMA Oncol, 1, 40.

6. George, J., Lim, J.S., Jang, S.J., Cun, Y., Ozretić, L., Kong, G., Leenders, F., Lu, X., Fernández-Cuesta, L., Bosco, G., et al. (2015) Comprehensive genomic profiles of small cell lung cancer. Nature, 524, 47–53.

7. Wheler, J.J., Janku, F., Naing, A., Li, Y., Stephen, B., Zinner, R., Subbiah, V., Fu, S., Karp, D., Falchook, G.S., et al. (2016) Cancer Therapy Directed by Comprehensive Genomic Profiling: A Single Center Study. Cancer Research, 76, 3690–3701.

8. Stubbington, M.J.T., Rozenblatt-Rosen, O., Regev, A. and Teichmann, S.A. (2017) Single-cell transcriptomics to explore the immune system in health and disease. Science, 358, 58–63.

9. Blom, K., Nygren, P., Alvarsson, J., Larsson, R. and Andersson, C.R. (2016) Ex Vivo Assessment of Drug Activity in Patient Tumor Cells as a Basis for Tailored Cancer Therapy. J Lab Autom, 21, 178– 187.

10. Lee, S.H.R., Yang, W., Gocho, Y., John, A., Rowland, L., Smart, B., Williams, H., Maxwell, D., Hunt, J., Yang, W., et al. (2023) Pharmacotypes across the genomic landscape of pediatric acute lymphoblastic leukemia and impact on treatment response. Nat Med, 29, 170–179.

11. Gawad, C., Koh, W. and Quake, S.R. (2014) Dissecting the clonal origins of childhood acute lymphoblastic leukemia by single-cell genomics. Proc. Natl. Acad. Sci. U.S.A., 111, 17947– 17952.

12. Zhang, Y., Wang, S., Zhang, J., Liu, C., Li, X., Guo, W., Duan, Y., Chen, X., Zong, S., Zheng, J., et al. (2022) Elucidating minimal residual disease of paediatric B-cell acute lymphoblastic leukaemia by single-cell analysis. Nat Cell Biol, 24, 242–252.

13. Van de Sande, B., Lee, J.S., Mutasa-Gottgens, E., Naughton, B., Bacon, W., Manning, J., Wang, Y., Pollard, J., Mendez, M., Hill, J., et al. (2023) Applications of single-cell RNA sequencing in drug discovery and development. Nat Rev Drug Discov, 22, 496–520.

14. Lim, J., Chin, V., Fairfax, K., Moutinho, C., Suan, D., Ji, H. and Powell, J.E. (2023) Transitioning single-cell genomics into the clinic. Nat Rev Genet, 10.1038/s41576-023-00613-w.

15. Bock, C., Boutros, M., Camp, J.G., Clarke, L., Clevers, H., Knoblich, J.A., Liberali, P., Regev, A., Rios, A.C., Stegle, O., et al. (2021) The Organoid Cell Atlas. Nat Biotechnol, 39, 13–17.

16. Mereu, E., Lafzi, A., Moutinho, C., Ziegenhain, C., McCarthy, D.J., Álvarez-Varela, A., Batlle, E., Sagar, null, Grün, D., Lau, J.K., et al. (2020) Benchmarking single-cell RNA-sequencing protocols for cell atlas projects. Nat Biotechnol, 38, 747–755.

17. Lindhagen, E., Nygren, P. and Larsson, R. (2008) The fluorometric microculture cytotoxicity assay. Nat Protoc, 3, 1364–1369.

18. Frost, B.-M., Forestier, E., Gustafsson, G., Nygren, P., Hellebostad, M., Jonsson, O.G., Kanerva, J., Schmiegelow, K., Larsson, R., Lönnerholm, G., et al. (2004) Translocation t(12;21) is related to in vitro cellular drug sensitivity to doxorubicin and etoposide in childhood acute lymphoblastic leukemia. Blood, 104, 2452–2457.

19. Lönnerholm, G., Thörn, I., Sundström, C., Frost, B.-M., Flaegstad, T., Heyman, M., Jonsson, O.G., Harila-Saari, A., Madsen, H.O., Porwit, A., et al. (2011) In vitro cellular drug resistance adds prognostic information to other known risk-factors in childhood acute lymphoblastic leukemia. Leukemia Research, 35, 472–478.

20. McGinnis, C.S., Patterson, D.M., Winkler, J., Conrad, D.N., Hein, M.Y., Srivastava, V., Hu, J.L., Murrow, L.M., Weissman, J.S., Werb, Z., et al. (2019) MULTI-seq: sample multiplexing for single-cell RNA sequencing using lipid-tagged indices. Nat Methods, 16, 619–626.

21. Hao, Y., Hao, S., Andersen-Nissen, E., Mauck, W.M., Zheng, S., Butler, A., Lee, M.J., Wilk, A.J., Darby, C., Zager, M., et al. (2021) Integrated analysis of multimodal single-cell data. Cell, 184, 3573–3587.e29.

22. Lun, A.T.L., participants in the 1st Human Cell Atlas Jamboree, Riesenfeld, S., Andrews, T., Dao, T.P., Gomes, T. and Marioni, J.C. (2019) EmptyDrops: distinguishing cells from empty droplets in droplet-based single-cell RNA sequencing data. Genome Biol, 20, 63.

23. Stuart, T., Butler, A., Hoffman, P., Hafemeister, C., Papalexi, E., Mauck, W.M., Hao, Y., Stoeckius, M., Smibert, P. and Satija, R. (2019) Comprehensive Integration of Single-Cell Data. Cell, 177, 1888–1902.e21.

24. Choudhary, S. and Satija, R. (2022) Comparison and evaluation of statistical error models for scRNA-seq. Genome Biol, 23, 27.

25. Ziegenhain, C., Vieth, B., Parekh, S., Reinius, B., Guillaumet-Adkins, A., Smets, M., Leonhardt, H., Heyn, H., Hellmann, I. and Enard, W. (2017) Comparative Analysis of Single-Cell RNA Sequencing Methods. Mol Cell, 65, 631–643.e4.

26. Finak, G., McDavid, A., Yajima, M., Deng, J., Gersuk, V., Shalek, A.K., Slichter, C.K., Miller, H.W., McElrath, M.J., Prlic, M., et al. (2015) MAST: a flexible statistical framework for assessing transcriptional changes and characterizing heterogeneity in single-cell RNA sequencing data. Genome Biol, 16, 278.

27. Wu, T., Hu, E., Xu, S., Chen, M., Guo, P., Dai, Z., Feng, T., Zhou, L., Tang, W., Zhan, L., et al. (2021) clusterProfiler 4.0: A universal enrichment tool for interpreting omics data. The Innovation, 2, 100141.

28. Yu, G., Wang, L.-G., Han, Y. and He, Q.-Y. (2012) clusterProfiler: an R Package for Comparing Biological Themes Among Gene Clusters. OMICS: A Journal of Integrative Biology, 16, 284– 287.

29. Subramanian, A., Tamayo, P., Mootha, V.K., Mukherjee, S., Ebert, B.L., Gillette, M.A., Paulovich, A., Pomeroy, S.L., Golub, T.R., Lander, E.S., et al. (2005) Gene set enrichment analysis: A knowledge-based approach for interpreting genome-wide expression profiles. Proc. Natl. Acad. Sci. U.S.A., 102, 15545–15550.

30. Jones, P.A. and Taylor, S.M. (1980) Cellular differentiation, cytidine analogs and DNA methylation. Cell, 20, 85–93.

31. Stresemann, C. and Lyko, F. (2008) Modes of action of the DNA methyltransferase inhibitors azacytidine and decitabine. Int. J. Cancer, 123, 8–13.

32. Rosenfeld, C., Goutner, A., Choquet, C., Venuat, A.M., Kayibanda, B., Pico, J.L. and Greaves, M.F. (1977) Phenotypic characterisation of a unique non-T, non-B acute lymphoblastic leukaemia cell line. Nature, 267, 841–843.

33. Bachmann, P.S., Gorman, R., Papa, R.A., Bardell, J.E., Ford, J., Kees, U.R., Marshall, G.M. and Lock, R.B. (2007) Divergent mechanisms of glucocorticoid resistance in experimental models of pediatric acute lymphoblastic leukemia. Cancer Res, 67, 4482–4490.

34. Mehtonen, J., Teppo, S., Lahnalampi, M., Kokko, A., Kaukonen, R., Oksa, L., Bouvy-Liivrand, M., Malyukova, A., Mäkinen, A., Laukkanen, S., et al. (2020) Single cell characterization of B-lymphoid differentiation and leukemic cell states during chemotherapy in ETV6-RUNX1-positive pediatric leukemia identifies drug-targetable transcription factor activities. Genome Med, 12, 99.

35. Dekker, L., Calkoen, F.G., Jiang, Y., Blok, H., Veldkamp, S.R., De Koning, C., Spoon, M., Admiraal, R., Hoogerbrugge, P., Vormoor, B., et al. (2022) Fludarabine exposure predicts outcome after CD19 CAR T-cell therapy in children and young adults with acute leukemia. Blood Advances, 6, 1969–1976.

36. Rosenberg, A.B., Roco, C.M., Muscat, R.A., Kuchina, A., Sample, P., Yao, Z., Graybuck, L.T., Peeler, D.J., Mukherjee, S., Chen, W., et al. (2018) Single-cell profiling of the developing mouse brain and spinal cord with split-pool barcoding. Science, 360, 176–182.

37. Ilicic, T., Kim, J.K., Kolodziejczyk, A.A., Bagger, F.O., McCarthy, D.J., Marioni, J.C. and Teichmann, S.A. (2016) Classification of low quality cells from single-cell RNA-seq data. Genome Biol, 17, 29.

38. Subramanian, A., Alperovich, M., Yang, Y. and Li, B. (2022) Biology-inspired data-driven quality control for scientific discovery in single-cell transcriptomics. Genome Biol, 23, 267.

39. Wang, X., He, Y., Zhang, Q., Ren, X. and Zhang, Z. (2021) Direct Comparative Analyses of 10X Genomics Chromium and Smart-seq2. Genomics Proteomics Bioinformatics, 19, 253–266.

40. Szklarczyk, D., Gable, A.L., Lyon, D., Junge, A., Wyder, S., Huerta-Cepas, J., Simonovic, M., Doncheva, N.T., Morris, J.H., Bork, P., et al. (2019) STRING v11: protein-protein association networks with increased coverage, supporting functional discovery in genome-wide experimental datasets. Nucleic Acids Res, 47, D607–D613.

41. Gutiérrez-Franco, A., Ake, F., Hassan, M.N., Cayuela, N.C., Mularoni, L. and Plass, M. (2023) Methanol fixation is the method of choice for droplet-based single-cell transcriptomics of neural cells. Commun Biol, 6, 522.

42. Mylka, V., Matetovici, I., Poovathingal, S., Aerts, J., Vandamme, N., Seurinck, R., Verstaen, K., Hulselmans, G., Van den Hoecke, S., Scheyltjens, I., et al. (2022) Comparative analysis of antibody- and lipid-based multiplexing methods for single-cell RNA-seq. Genome Biol, 23, 55.

43. Brown, D.V., Anttila, C.J.A., Ling, L., Grave, P., Baldwin, T.M., Munnings, R., Farchione, A.J., Bryant, V.L., Dunstone, A., Biben, C., et al. (2023) A Risk-reward Examination of Sample Multiplexing Reagents for Single Cell RNA-Seq Genomics.

44. Engeland, K. (2022) Cell cycle regulation: p53-p21-RB signaling. Cell Death Differ, 29, 946–960.

45. Huang, S., Shu, L., Dilling, M.B., Easton, J., Harwood, F.C., Ichijo, H. and Houghton, P.J. (2003) Sustained Activation of the JNK Cascade and Rapamycin-Induced Apoptosis Are Suppressed by p53/p21Cip1. Molecular Cell, 11, 1491–1501.

46. Elhasasna, H., Khan, R., Bhanumathy, K.K., Vizeacoumar, F.S., Walke, P., Bautista, M., Dahiya, D.K., Maranda, V., Patel, H., Balagopal, A., et al. (2022) A Drug Repurposing Screen Identifies Fludarabine Phosphate as a Potential Therapeutic Agent for N-MYC Overexpressing Neuroendocrine Prostate Cancers. Cells, 11, 2246.

47. Ghandi, M., Huang, F.W., Jané-Valbuena, J., Kryukov, G.V., Lo, C.C., McDonald, E.R., Barretina, J., Gelfand, E.T., Bielski, C.M., Li, H., et al. (2019) Next-generation characterization of the Cancer Cell Line Encyclopedia. Nature, 569, 503–508.

48. Shin, D., Lee, W., Lee, J.H. and Bang, D. (2019) Multiplexed single-cell RNA-seq via transient barcoding for simultaneous expression profiling of various drug perturbations. Sci. Adv., 5, eaav2249.

49. Stoeckius, M., Hafemeister, C., Stephenson, W., Houck-Loomis, B., Chattopadhyay, P.K., Swerdlow, H., Satija, R. and Smibert, P. (2017) Simultaneous epitope and transcriptome measurement in single cells. Nat Methods, 14, 865–868.

