## Supplementary Figures for "COMPARISON OF HIGH-THROUGHPUT SINGLE-CELL RNA-SEQ METHODS FOR EX VIVO DRUG SCREENING"

Figure S1

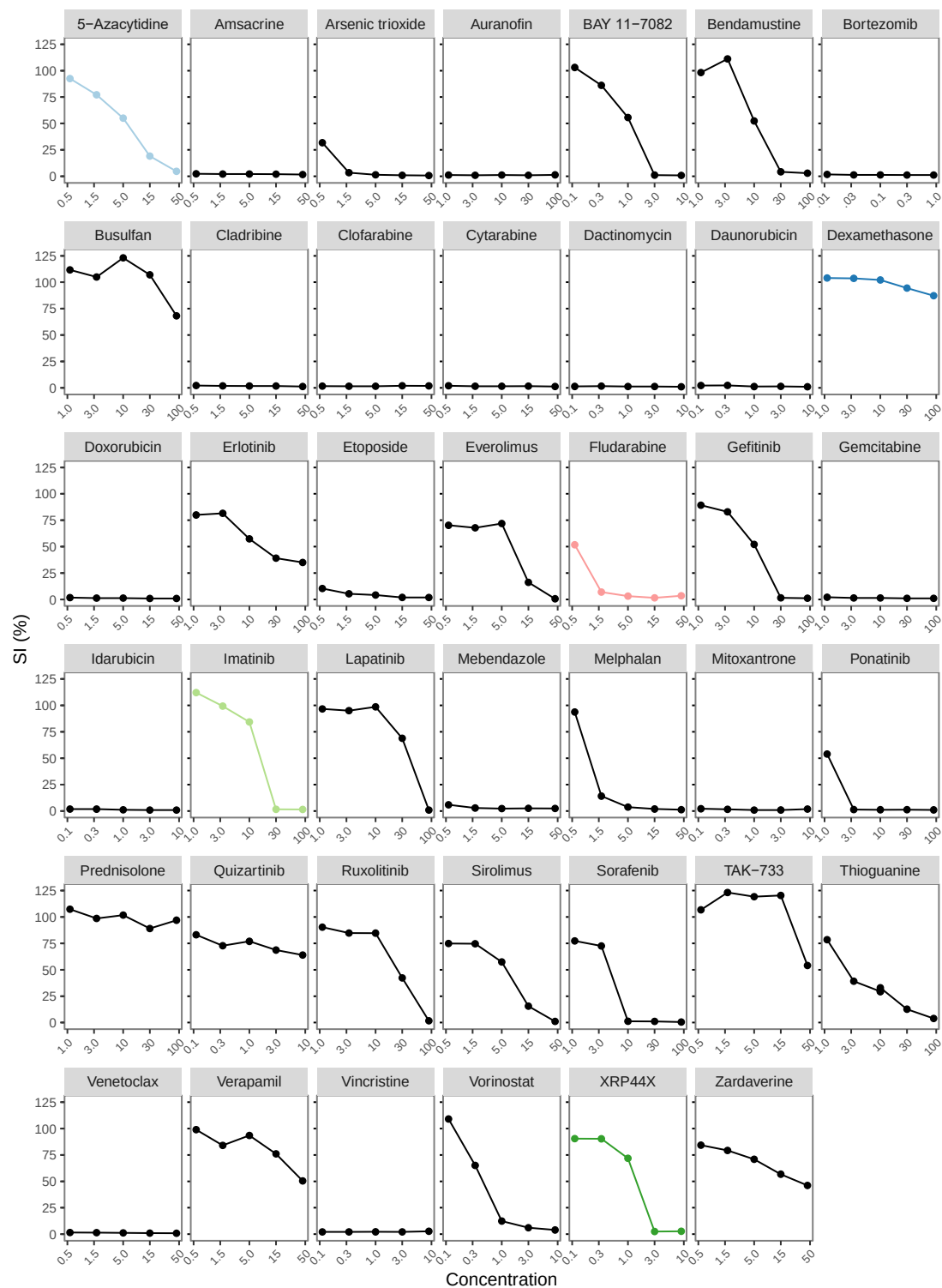

**Supplementary Figure S1.** Survival index of Reh cells exposed to 72 hours treatment of cytostatic treatment at the concentrations as indicated in each panel. Each dot represents the mean measured from two replicates. The cells surviving the following treatments were selected for single-cell RNA-seq: 5-Azacytidine (light blue), Dexamethasone (dark blue), Fludarabine (pink), Imatinib (light green), and XRP44X (dark green).

Figure S2

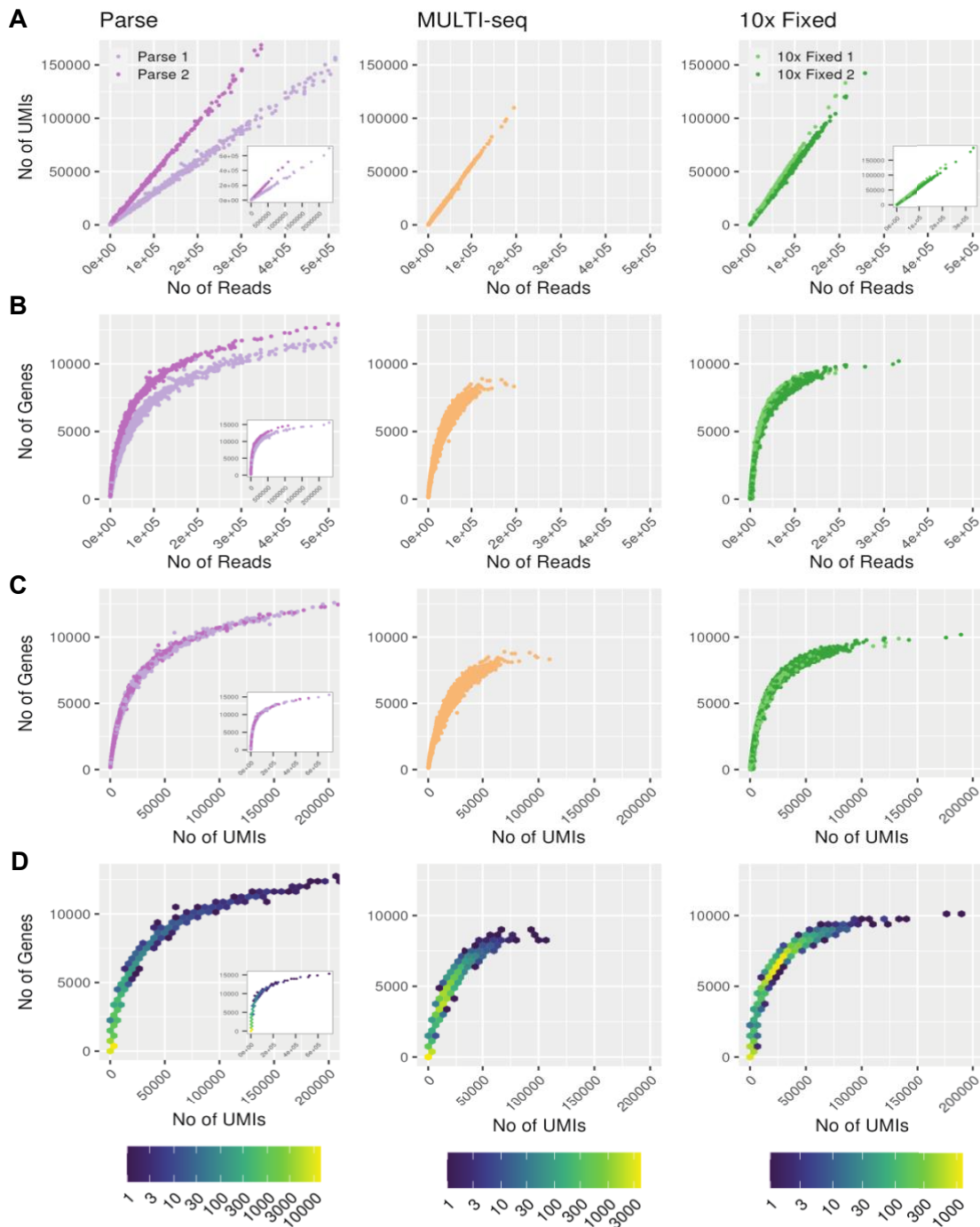

**Supplementary Figure S2.** Comparisons of sequencing depth, UMIs and number of genes detected prior to cell filtering for each scRNA-seq library method. (A) The number of reads (x-axis) plotted against the number of UMIs (y-axis). Note that each sub-library has a linear correlation, while in the sub-libraries with different read depth the correlation factor is not equal. The inset for the Parse data shows the full range of barcodes. (B) The number of genes detected plotted against number of reads for each cell-barcode. (C) The number of genes detected plotted against number of UMIs for each cell-barcode. (D) The number of genes detected plotted against the number of UMIs detected per cell-barcode visualized as hexagonal binning plot, the color indicates the number of cell-barcodes contributing to each hexagonal bin according to each scale below each panel.

Figure S3

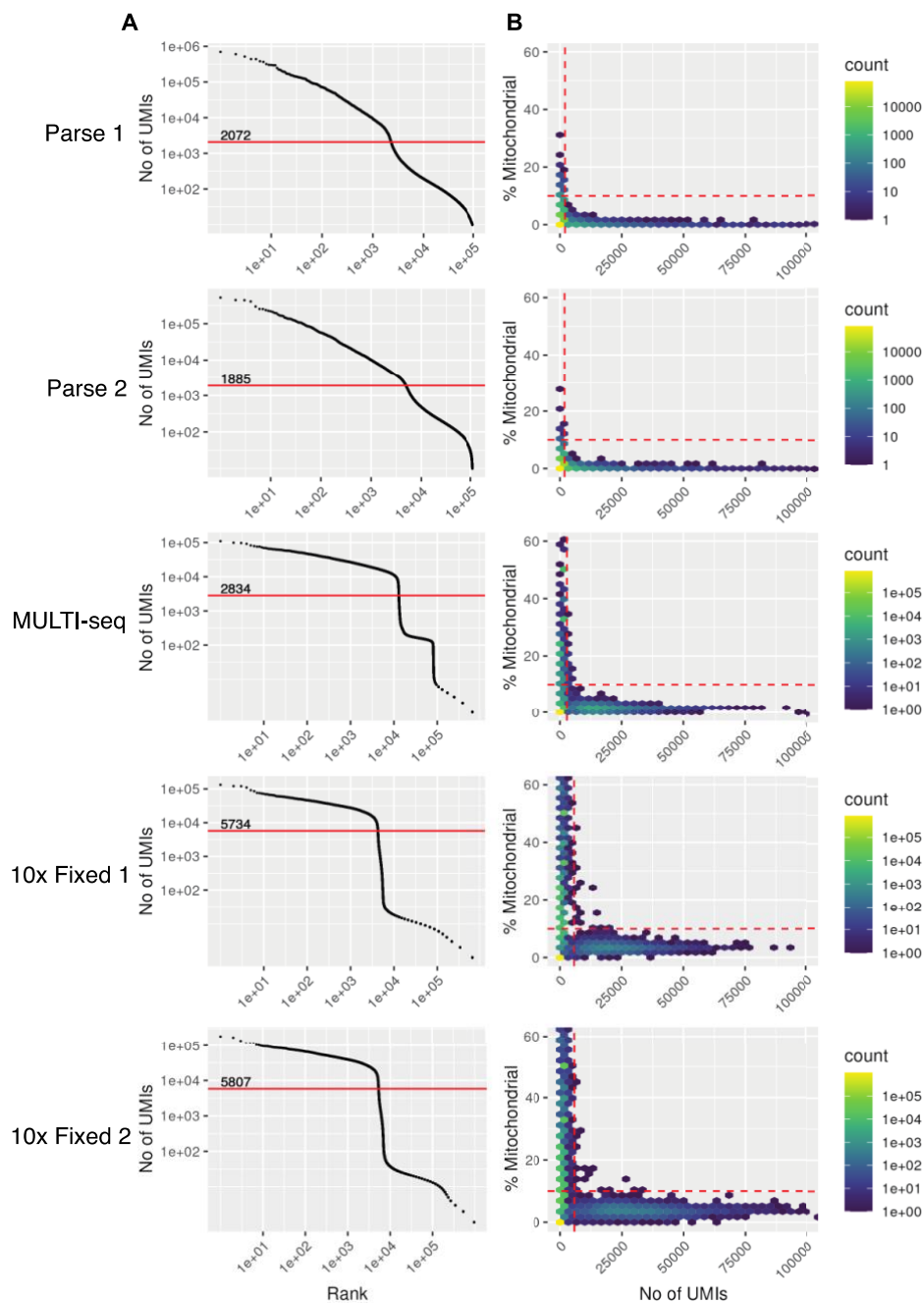

**Supplementary Figure S3.** Filtering selection for removing non-cell UMIs. (A) Knee bar plots for each sequencing library, the y-axis indicates the number of UMIs and the x-axis the rank of each index. The library (sub-library) is indicated to the left of the panel. The red line indicates the inflection point, only barcodes with detected UMIs above the given number are considered to be cells used for downstream analyses. (B) Histogram of the percent of reads mapping to mitochondrial genes (y-axis) and the number of UMIs (x-axis), visualized as hexagonal binning plots. The color indicates the number of cells contributing to each hexagonal bin according to the scale to the right of each panel. The dashed red vertical line indicates the cut-off of the inflection point from the corresponding plot to the left (panel A). The dashed red horizontal line indicates the cut-off for filtering out cells with >10% sequencing reads that map to mitochondrial genes.

Figure S4

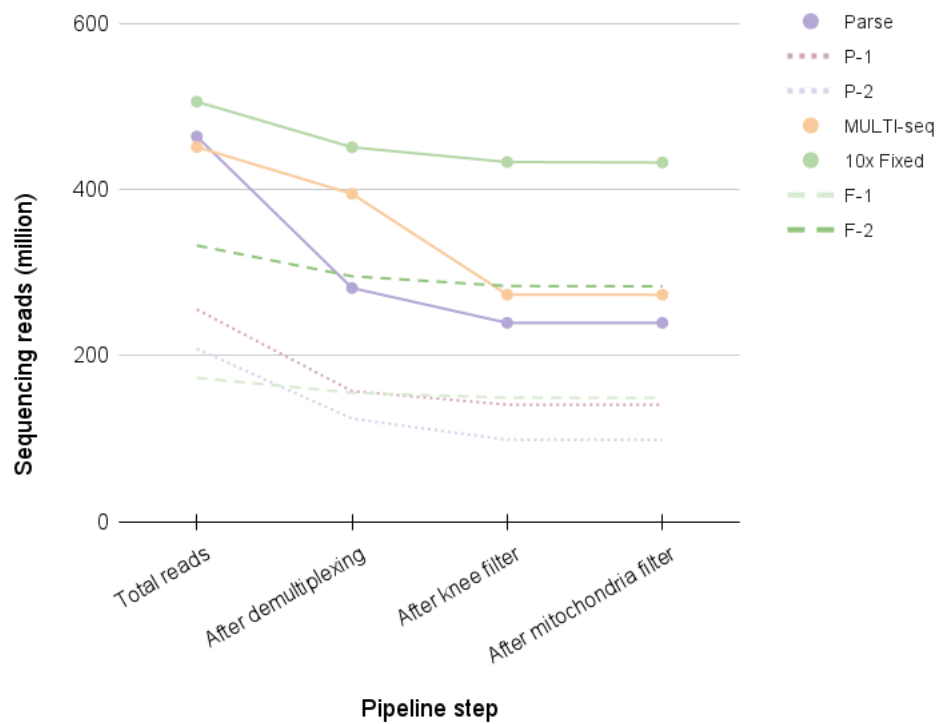

**Supplementary Figure S4.** Reads remaining during the data processing pipeline. The reads remaining after each step in the pipeline are indicated by method (Parse, MULTI-seq, 10x Fixed) and sub-library (P-1, P-2, F-1 and F-2) according to the legend to right of the panel.

Figure S5

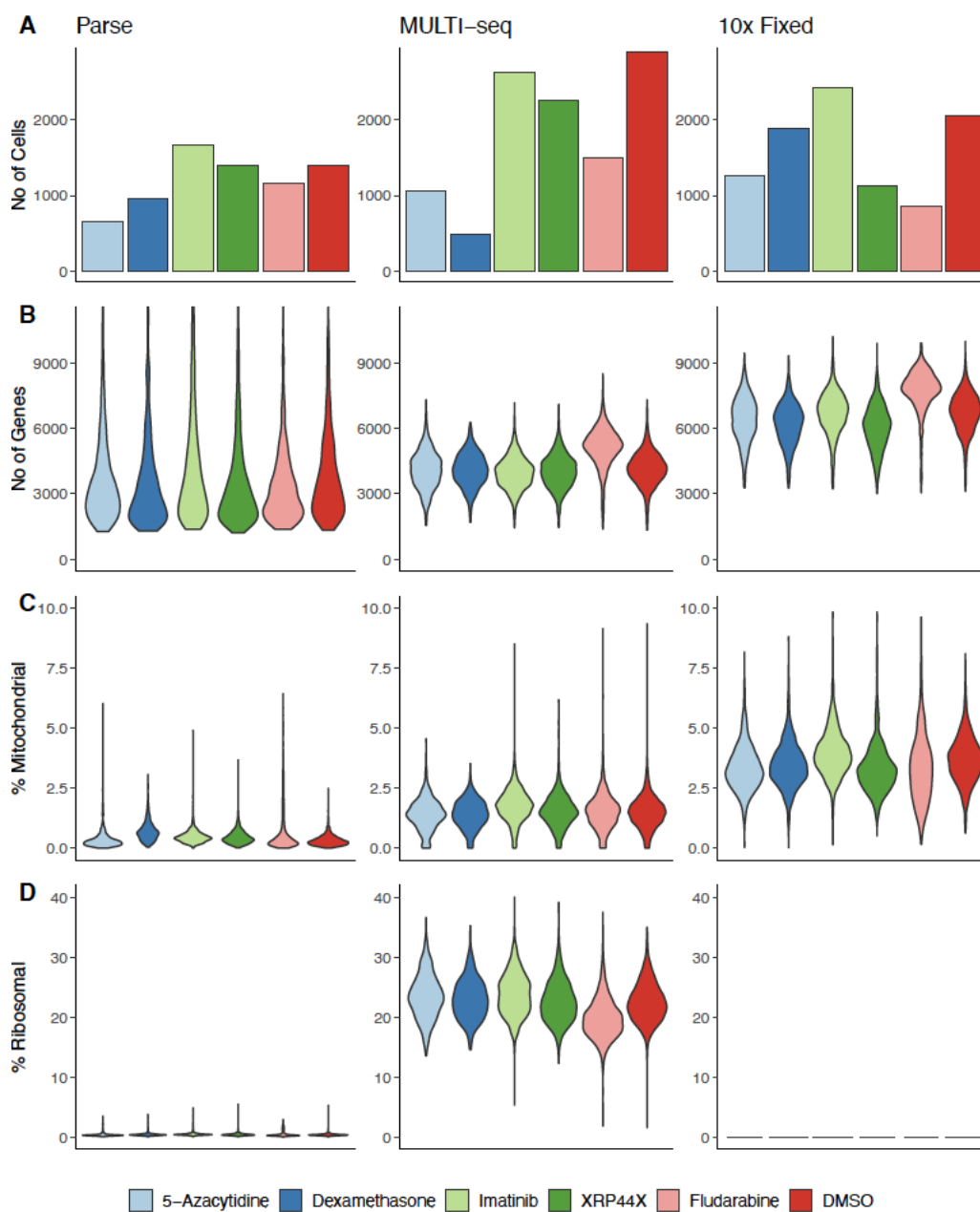

**Supplementary Figure S5.** Quality metrics of single cell RNA sequencing (scRNA-seq) data generated from three different library preparation methods (Parse, MULTI-seq, 10x Fixed) separated by drug condition (barcode). The color key for the drug conditions is indicated at the bottom of the figure. (A) The number of cells detected by each scRNA-seq library preparation method and condition presented as bar plots. (B) The number of genes detected by each scRNA-seq library preparation method and condition presented as violin plots. (C) The percentage of reads that map to mitochondrial genes by scRNA-seq library preparation method and condition presented as violin plots. (D) The percentage reads that map to ribosomal genes by scRNA-seq library preparation method and condition presented as violin plots.

Figure S6

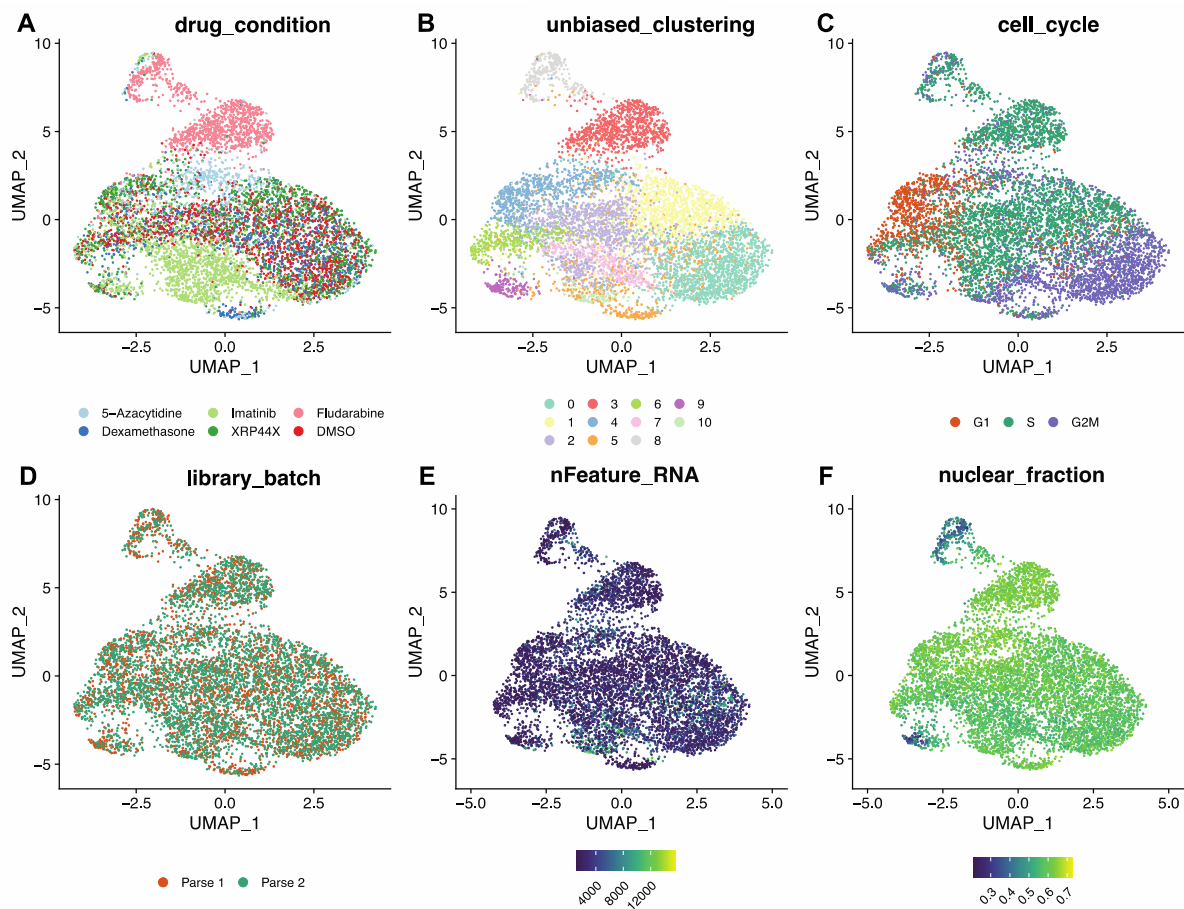

**Supplementary Figure S6.** UMAP visualization of Parse single-cell RNA-seq data. The cells are color-coded to depict various biological attributes: (A) Drug condition/barcode, (B) Unbiased clustering using FindClusters in the R package "Seurat", (C) Cell cycle estimation, (D) Sub-library, (E) Number of genes detected, and (F) Fraction of nuclear reads, calculated as $N_{\text{intronic reads}} / (N_{\text{intronic reads}} + N_{\text{exonic reads}})$ using the R package "DropletQC".

Figure S7

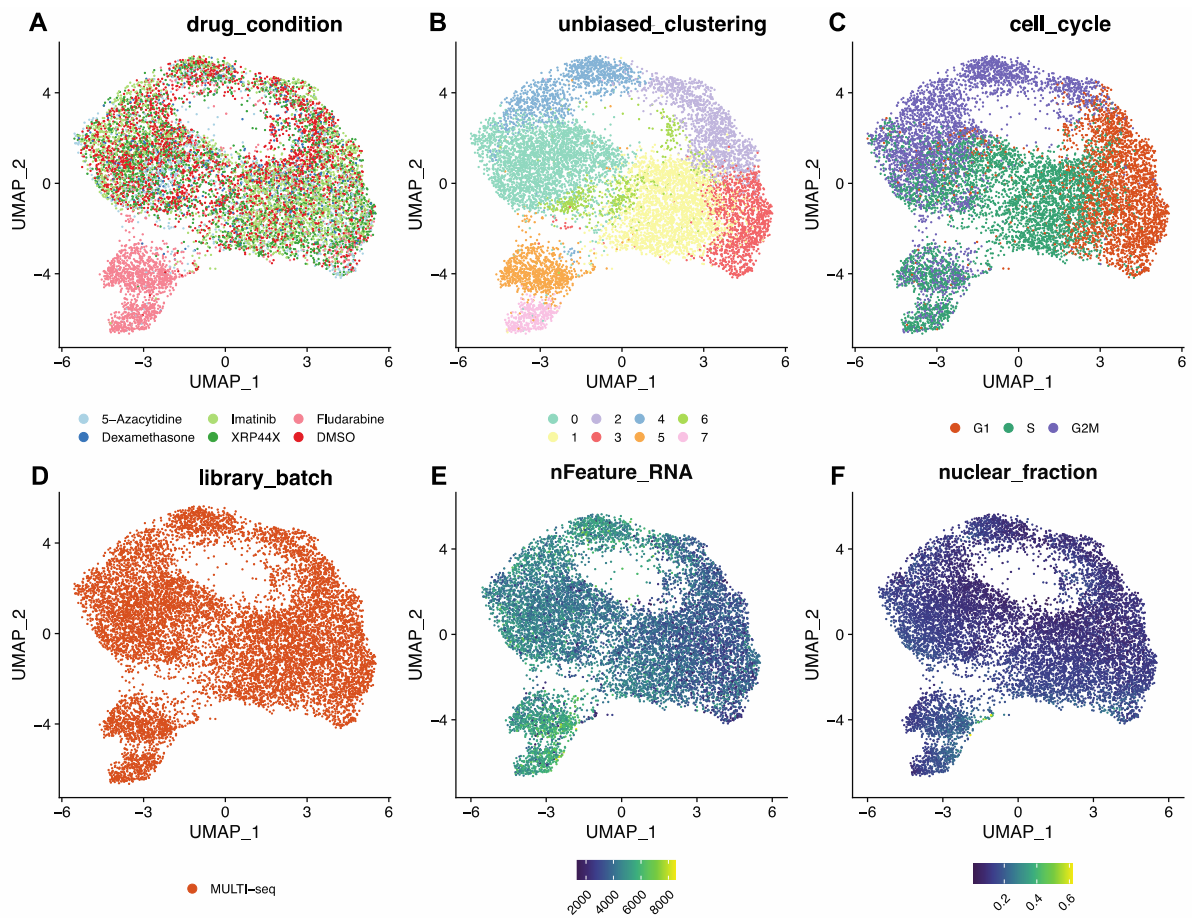

**Supplementary Figure S7.** UMAP visualization of MULTI-seq single-cell RNA-seq data. The cells are color-coded to depict various biological attributes: (A) Drug condition/barcode, (B) Unbiased clustering using FindClusters in the R package "Seurat", (C) Cell cycle estimation, (D) library batch (for MULTI-seq there was only one batch), (E) Number of genes detected, and (F) Fraction of nuclear reads, calculated as $N_{\text{intronic reads}} / (N_{\text{intronic reads}} + N_{\text{exonic reads}})$ using the R package "DropletQC".

Figure S8

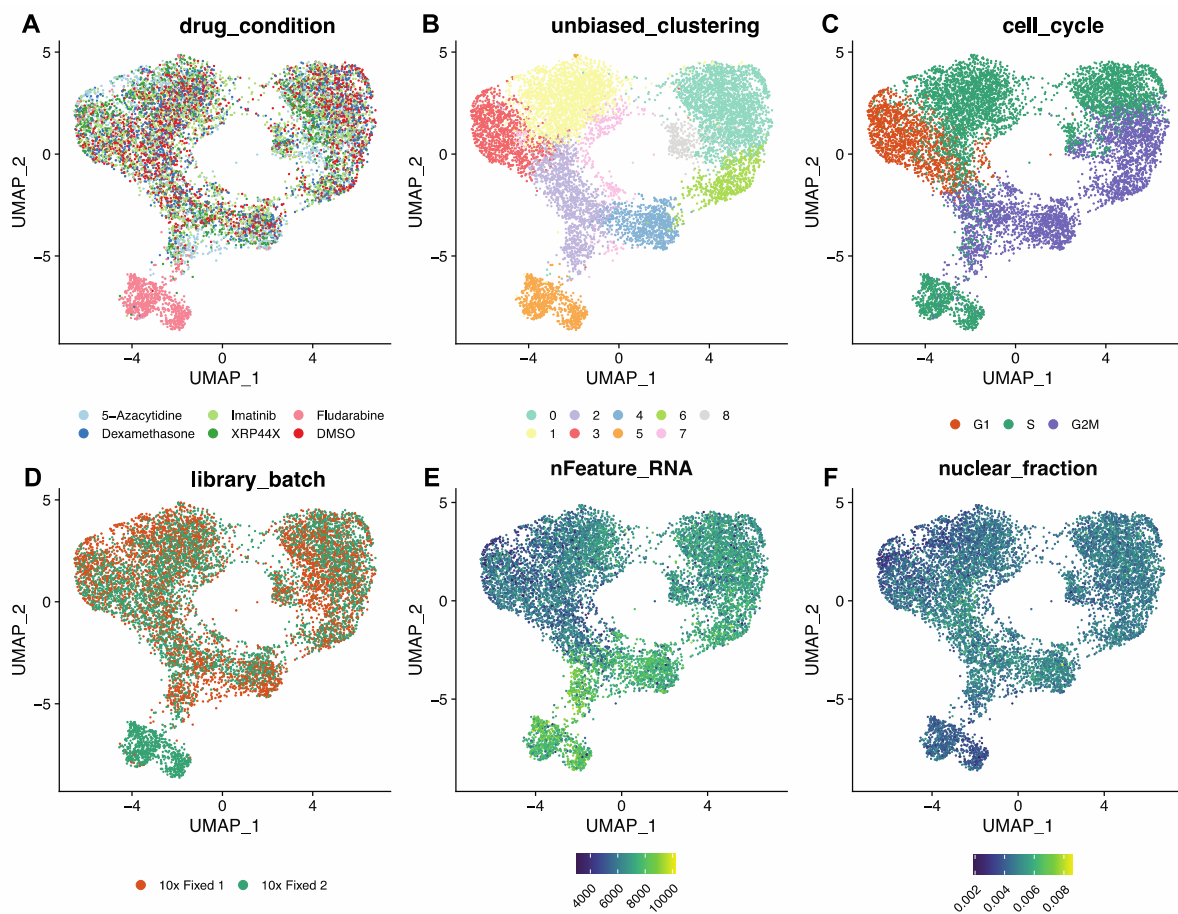

**Supplementary Figure S8.** UMAP visualization of 10x Fixed single-cell RNA-seq data. The cells are color-coded to depict various biological attributes: (A) Drug condition/barcode, (B) Unbiased clustering using FindClusters in the R package "Seurat", (C) Cell cycle estimation, (D) Library preparation batch, (E) Number of genes detected, and (F) Fraction of nuclear reads, calculated as $N_{\text{intronic reads}} / (N_{\text{intronic reads}} + N_{\text{exonic reads}})$ using the R package "DropletQC".

Figure S9

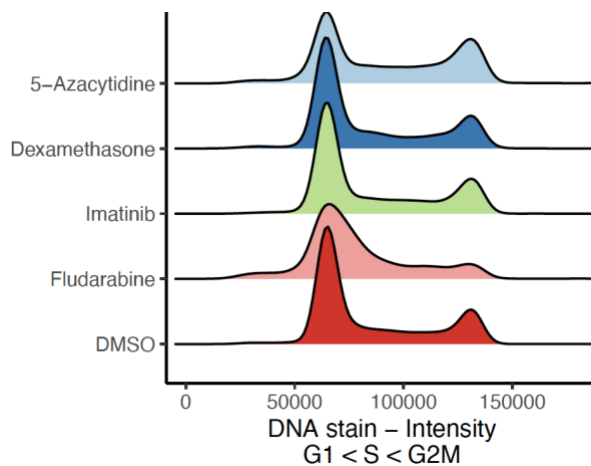

**Supplementary Figure S9.** Cell cycle states after 72 hours of treatment. Ridge plot with population histogram with the DNA content measured by DAPI intensity (x-axis), the density of cells for each treatment is represented on the y-axis.

Figure S10

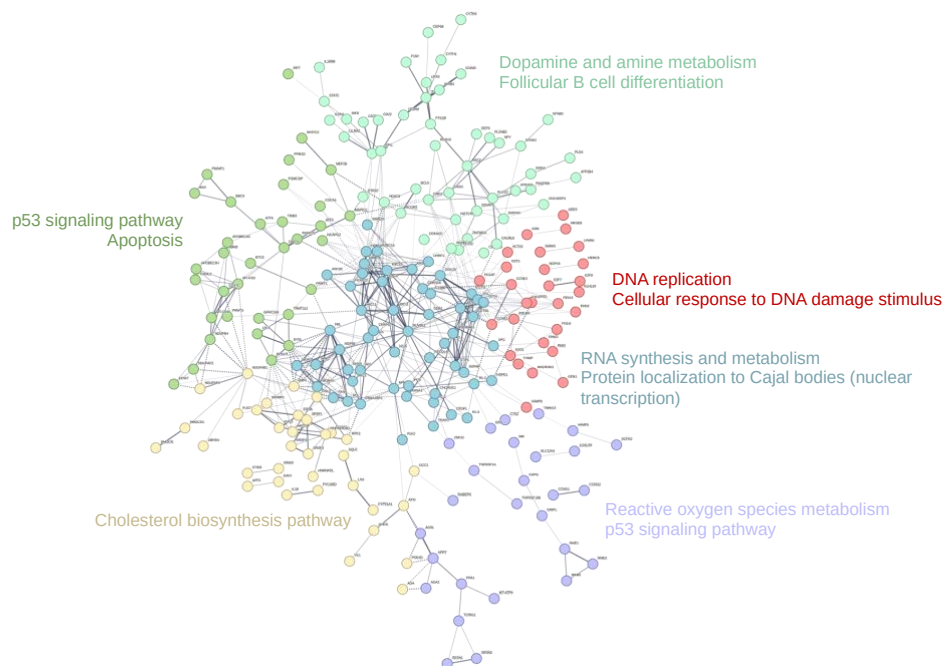

**Supplementary Figure S10.** STRING network of 441 differentially expressed genes detected by any scRNAseq method in fludarabine-treated vs control cells. Protein interactions have been selected on co-expression and experimental evidence parameters, and further categorized into six distinct functional groups by K-means clustering. Proteins without a functional or physical association to any other protein in the network have been excluded from the interaction plot.
